# Cloning and characterization of a novel maize leaf area modifier and its effects across elite germplasm

**DOI:** 10.64898/2026.05.15.725441

**Authors:** Matthew J. Runyon, Marlee R. Labroo, Miriam I. Arend, Michael J. Scanlon, Anthony J. Studer

## Abstract

Plant architecture is a crucial component of maize productivity. Tailoring architectural component traits like leaf area and angle can increase productivity by promoting deeper light penetration into the canopy and better resource utilization. Novel genetic variants can increase the rate of gain for optimized plant architecture. Here, we map a moderate-effect mutation denoted *reduced leaf area1* (*rdla1*) to the *RAGGED5* (*RGD5*) locus and characterize it as a transposon insertion allele. Mutant leaf area reductions were most extreme in mid-upper canopy positions. Photosynthetic gas exchange rates were not significantly impacted in *rdla1* relative to wild-type, indicating that mutant leaf structure, but not function, is altered. Functional annotations of *RDLA1* were supported by metabolite profiles suggesting a role in cuticular wax biosynthesis. Introgression of the *rdla1* allele into 27 commercially relevant genetic backgrounds identified differences in effect size across genotypes, revealing modifier effects that could serve as targets for modulating plant architecture.

## Introduction

Maize (*Zea mays* L.) breeding programs heavily focus on selecting traits that improve grain yield^1^. Yield is recognized as the product of solar radiation with interception, conversion, and partitioning efficiencies^2^. Historical gains in grain yield have resulted mostly from improved light interception and partitioning efficiencies, which are arguably approaching their maxima^3^. Increases in light interception have been accomplished through increased leaf area index (LAI), driven across breeding eras by increased planting density^4^. In the United States, planting density has increased by approximately 600 plants per hectare per year from 1960 through the present date^5^. Throughout contemporary maize breeding in North America, there has been a strong, positive correlation between planting density and yield improvement^6,7^. Attempts to quantify the impact of density on maize grain yield suggest up to 17% of yield gains may be attributed to increased planting density^8^.

Increasing planting density improves total grain yield of a defined area, despite a reduction in the grain harvested from each plant due to resource competition. Up to a threshold, this loss in individual plant yield is more than offset simply by having a greater gross number of plants in a given space^9^. A maximum threshold for planting density exists based on available environmental resources, and further increasing the number of plants beyond this threshold will decrease grain yield^10^. Reduced interplant competition at high densities may be achieved with narrower inter-row spacing coupled with wider intra-plant spacing within rows^9,11,12^. Beyond management strategies, breeding programs have also selected loci conferring adaptation to high planting density^13,14^. However, genetic traits governing source-sink relations, photosynthetic capacity, stress tolerance, growth rate, and canopy architecture all interact with management practices and may synergistically or antagonistically impact density tolerance^15^. A primary component trait that controls canopy architecture and photosynthetic capacity is leaf area. Plants with smaller leaves at the ear-bearing node have a competitive yield advantage over plants with larger leaves at that position when selected out of a common genetic background^16^. However, assessing the direct effect of smaller leaves on yield across different breeding eras and genetic backgrounds is confounded by concurrent selection of other agronomically valuable traits in tandem^17^.

Crop canopy traits are primarily regulated by quantitative, small-effect loci, and these factors have likely been subject to indirect selection within breeding programs^18,19,20^. Improvements to canopy architecture using qualitative loci have been achieved with limited success. Early work demonstrated that hybrids of *liguleless1* (*lg1*) and *liguleless2* (*lg2*) mutant introgression lines conferred more upright leaves that were directly correlated to elevated yields at increasing plant density^21^. While introgression of these null alleles did not take place at scale in breeding pipelines, selection of genotypes with altered regulation of these genes was a significant driver of improved adaptation to elevated planting density among contemporary germplasm^18^.

Qualitative loci modifying leaf area itself in maize have been previously described, yet most natural alleles cause phenotypes that are too severe to be used in agronomic systems. For example, homozygous *Liguless narrow-Reference* (*Lgn-R*) mutants are permanently juvenile, while heterozygous individuals confer a moderated effect of reduced blade width and length with penetrance heavily impacted by genetic background^22,23^. The *narrow sheath* (*ns*) mutant results in a deletion of the marginal leaf domain that extends from the mid-length of the blade to include the entire leaf sheath, a phenotype that is most readily observed in leaves at lower canopy positions^24^. The *ns* phenotype is conferred by loss-of-function mutations at the duplicate *ns1* and *ns2* loci, which encode redundant WOX3 transcription factors that recruit the marginal leaf domain from the shoot apical meristem (SAM)^25,26^. Likewise, *rough sheath2* (*rs2*), mutants are defective in the regulation of Class 1 *KNOX* genes and produce leaves with multiple midribs that are often bladeless in certain genetic backgrounds^27,28,29^. Pleiotropic effects of dwarfing and sinuous internodes were observed in *ns* mutants, while *rs2* mutants had twisted internodes and reduced floret numbers sometimes coupled with partial male sterility, limiting their practical utility^24,27^.

A single-locus mutation denoted *reduced leaf area1* (*rdla1*) confers a more moderate leaf area reduction than other known qualitative leaf area mutants. First identified in 1970, maize inbreds containing this recessive mutation displayed significant reductions in leaf area of the ear-bearing node relative to wild-type^30^. Hybrids exhibiting reduced ear leaf areas have previously shown a competitive advantage over hybrids with larger ear leaf areas when planted at high density^16^. As a single mutation with mild effect on leaf area, *rdla1* may serve as a breeding target for reduced leaf area while enabling the discovery of associated modifier loci.

Through a forward genetics approach, we characterized the *rdla1* phenotype and identified the causative mutation using traditional recombinant mapping and sequenced-based approaches. Introgression of the *rdla1* locus into a panel of 27 maize Expired Plant Variety Protection (ExPVP) lines, which are genotypes originating from commercial breeding programs, revealed a gradient of leaf area phenotypes across genetic backgrounds. Differential patterns in leaf and canopy structure conferred by genetic background effects interacting with the *rdla1* allele were observed. Collectively, these findings suggest that designing an idealized canopy structure for optimized productivity in modern agronomic conditions is possible through the introgression and modulation of genes like *rdla1*.

## Results

### Identification of the rdla1 Locus

To identify the mutation underlying the *rdla1* phenotype, both recombinant mapping and sequencing approaches were utilized. Initially, three rounds of mapping were iteratively conducted. Whole-genome shotgun sequencing of four Oh43 wild-type and four Oh43 BC_5_S_10_ individuals placed the genomic region of the *rdla1* locus on the long arm of Chromosome 4. Subsequent sequencing of 26 Oh43 *rdla1* BC_6_S_1_ and four wild-type individuals in addition to the eight BC_5_S_10_ individuals previously sequenced revealed that *rdla1*-associated sites clustered between 176.3Mb and 180.4Mb of Chromosome 4 in the B73_v5 reference genome. A final iteration used 182 Oh43 *rdla1* BC_6_S_1_ individuals in addition to the 26 BC_6_S_1_ individuals used in the second round of fine mapping. These combined 208 BC_6_S_1_ individuals were screened for informative recombination events using PCR-based markers in the identified 4.1Mb region. 17 such individuals were found to have recombination in the target interval and were bulk whole-genome sequenced. Among all rounds of fine-mapping, the peak of the *rdla1*-associated sites was found in a bin of ∼151kb corresponding to coordinates 4:176786643 to 4:176938025 in the B73_v5 reference genome.

Using the information from the bulk-segregant analysis to define the *rdla1* candidate region, we designed a series of 10 indel markers (M1 – M10; Supplementary Table 1) for marker-assisted backcrossing to introduce the mutant allele into an ExPVP germplasm panel (Figure 1; Supplementary Table 2). These markers spanned a region of ∼1.38Mb from coordinates 4:176530120 to 4:177912638 in the B73_v5 reference genome, containing 58 annotated genes. Using this marker set to track recombination events while backcrossing the mutation into B73 and 27 ExPVP maize lines from the BC_1_ to BC_5_S_1_ generations, we narrowed the candidate interval to the region between M4 and M5. An additional marker developed for detecting polymorphisms via Sanger sequencing (S1) narrowed the candidate region further to an ∼82kb region between S1 and M5 that contained five genes (Figure 1).

**Figure 1:**
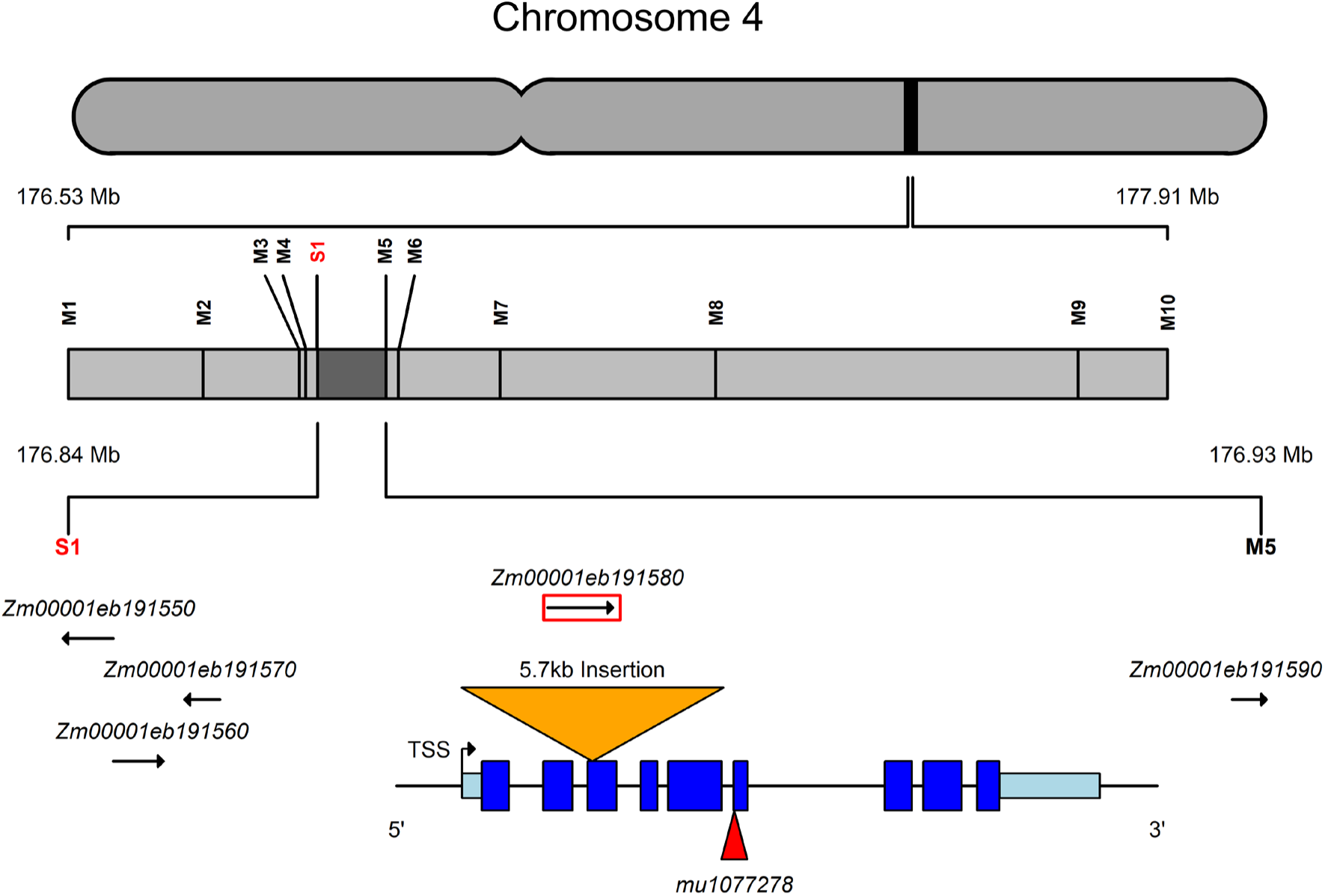
Genomic position of *rdla1* mutation on Chromosome 4. A candidate region spanning 176.53Mb – 177.91Mb was established via bulk segregant analysis. Indel markers used for mapping recombination events within the candidate region are denoted M1 – M10. The Sanger sequencing marker used to narrow the recombination interval is denoted S1. B73 gene IDs contained within the fine mapping region (176.84Mb – 176.93Mb) are shown, with the candidate gene outlined in red. Below the red outlined box, the gene model shows canonical exon structure of candidate gene *Zm00001eb191580*. Exons are shown in dark blue, while 5’ and 3’ UTRs are shown in light blue. The insertion positions of the *Magellan* element underlying *rdla1* (orange triangle) and *mu1077278* (red triangle) are shown.

To identify any large-scale structural differences within the candidate region, Oh43 *rdla1* BC_5_S_11_ genomic DNA underwent PacBio sequencing and was aligned to the Oh43 reference genome. The assembled sequences revealed an insertion of ∼5.7kb in *Zm00001eb191580* that was not present in wild-type (Figure 1). This insertion site was located after the 47^th^ base pair of Exon 6 and introduces a premature stop codon. This insertion sequence bears 99.95% identity to the *Magellan*-family retrotransposon AC194897-4313. *Zm00001eb191580* is annotated as an aldehyde decarbonylase with a predicted functional role in cuticular wax biosynthesis and underlies the *RAGGED5* (*RGD5*) locus.

### Complementation Test

To validate that the *Magellan* retrotransposon insertion in *Zm00001eb191580* is responsible for the *rdla1* phenotype, a genetic complementation test was performed. The UFMu-09400 stock contains a *Mu* transposable element located within the sixth exon of *Zm00001eb191580* (*mu1077278;* Figure 1). Plants heterozygous for *mu1077278* were crossed to wild-type individuals of PHKW3, an ExPVP inbred which shows striking visual reductions in leaf area when homozygous for the *rdla1* introgression. A heterozygous *mu1077278*/+ F_1_ plant was crossed with a homozygous *rdla1/rdla1* PHKW3 BC_5_S_3_ individual. Progeny from this cross were grown in the greenhouse, and genotyping confirmed that the progeny segregated in an expected 1:1 ratio of *rdla1/+* and *rdla1/ mu1077278*. Leaf area of the *rdla1/ mu1077278* individuals was significantly reduced relative to *rdla/+* individuals at the ear leaf (–26.95%) and at the third leaf above (–34.29%) and below (–22.63%) the ear-bearing node (n = 8, Welch’s *t*-test, *p <* 0.0001) (Figure 2). Leaf width and length reductions in *rdla1/ mu1077278* plants both contributed to this reduction in leaf area (Supplementary Figure 1). Because the *rdla1* allele failed to complement the insertion allele, we demonstrated that the retrotransposon insertion in *Zm00001eb191580* is causal for the *rdla1* phenotype.

**Figure 2:**
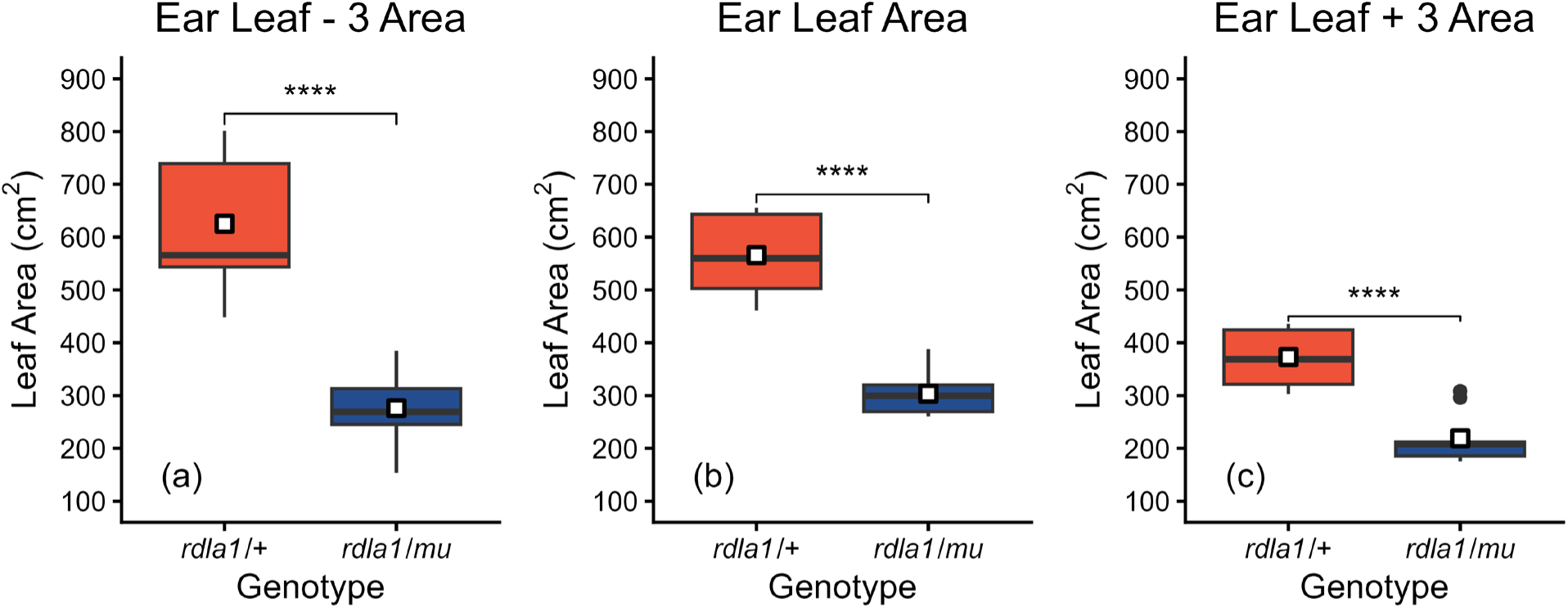
Absolute leaf areas of *rdla1/+* (orange) and *rdla1/mu1077278* (blue) plants at the third leaf below the ear-bearing node (a), the leaf of the ear-bearing node (b), and the third leaf above the ear bearing node (c) (n = 8 plants). The centerline indicates the median, while the white square indicates the mean. The box is bounded by the lower 25^th^ percentile and upper 75^th^ percentile. Whiskers extend to the largest and smallest values within 1.5× the interquartile range for the 75^th^ and 25^th^ percentiles, respectively, while points beyond whiskers represent values < 3× the interquartile ranges. Significant differences calculated using a two-tailed Welch’s *t*-test are denoted as follows: **** = p < 0.0001, *** = p < 0.001, ** = p < 0.01, * = p < 0.05, ns = nonsignificant.

### Phenotypic Characterization of rdla1 Across Developmental Time

To fully characterize the phenotypic effect of the *rdla1* mutation, the mutant allele was evaluated in B73 BC_5_S_2_ introgression lines. For advanced characterization of the *rdla1* mutation throughout developmental time, we measured the length, width, and area of each leaf from Leaf 2 through Leaf 20. While reduced leaf area was observed at all leaf positions, a significant reduction in area was found predominantly in the mid-upper canopy at Leaf 11 –19 positions (Figure 3a). Differences in both length and width contributed to this reduction in area (Supplementary Figures 2a-b). The magnitude of the leaf area reduction conferred by the *rdla1* mutation varied depending on leaf position. While the mid-upper canopy had the largest leaf areas in both wild-type and *rdla1* plants, leaves at positions Leaf 13 –18 also had the greatest reduction in area (a maximum reduction of –32.09% ± 1.50% at the Leaf 17 position; Figure 3b). In B73 *rdla1* and wild-type lines, the leaf of the ear-bearing node ranged from Leaf 15 (–30.11% ± 1.08% leaf area reduction) to Leaf 16 (–31.18% ± 0.85% leaf area reduction). Neither plant height nor the height of the ear-bearing node were significantly different between wild-type and *rdla1* plants (n = 4 blocks, Welch’s *t*-test, p > 0.05) (Supplementary Figure 3).

**Figure 3:**
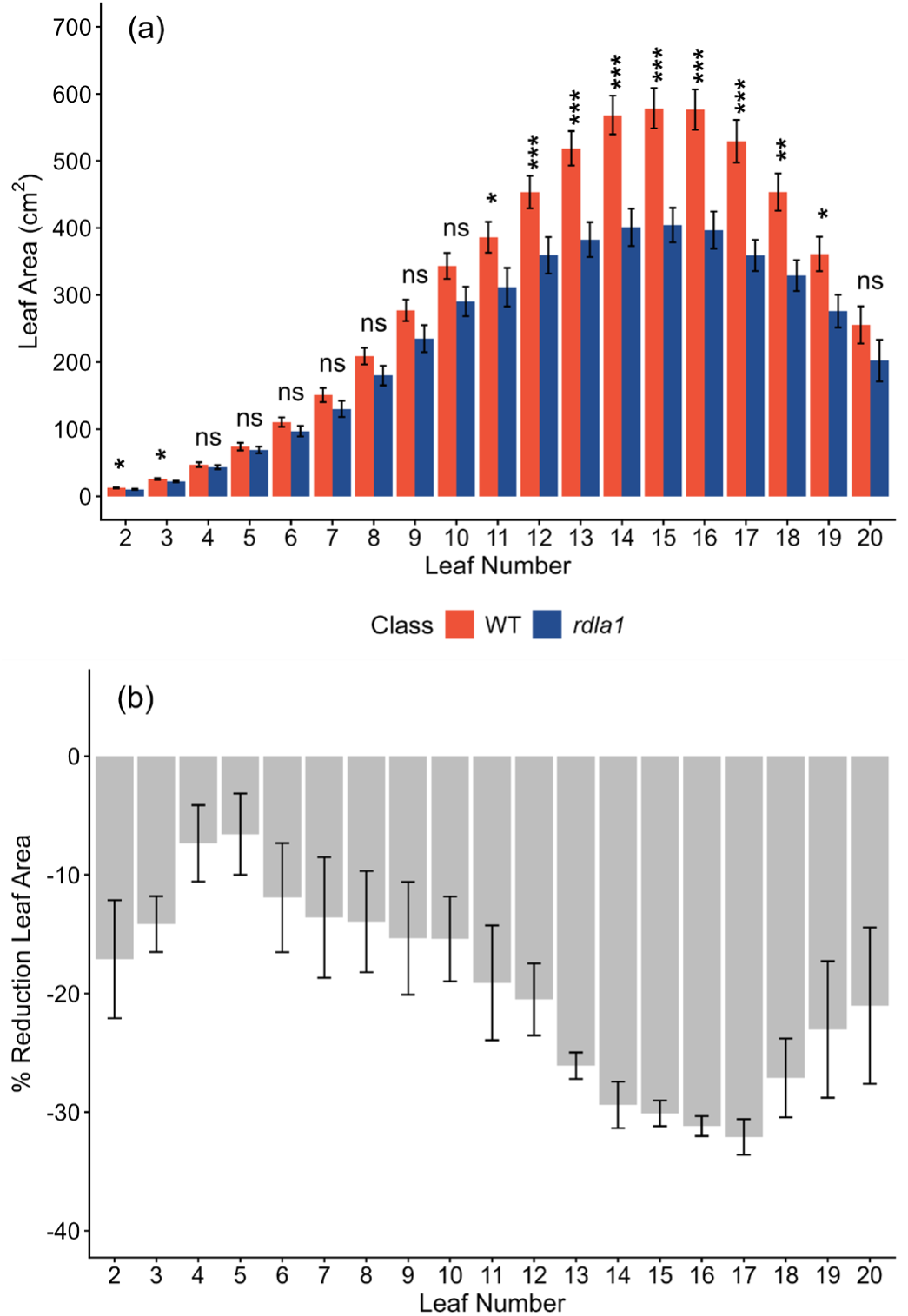
(a) The mean absolute leaf area of wild-type (orange) and *rdla1* (blue) at leaf positions 2 through 20. (b) The mean percentage reduction of *rdla1* leaves relative to wild-type at each leaf position. Bars denoting mean leaf area and percentage reductions are representative of plots (n = 4) ± standard error. Significant differences calculated using a two-tailed Welch’s *t*-test are denoted as follows: *** = p < 0.001, ** = p < 0.01, * = p < 0.05, ns = nonsignificant.

### Anatomical and Physiological Characterization

Anatomical and physiological characterization of the *rdla1* allele were completed in a B73 genetic background. Additionally, LH198 and 3IIH6 were selected from the panel of 27 ExPVP *rdla1* introgression lines for additional characterization due to known historical importance of these lines in commercial breeding programs. Transverse sectioning of Leaf 2 in B73 showed anatomy and physiology that was equivalent to wild-type, with all leaf domains present in both genotypic classes (Supplementary Figure 4a-f). Likewise, no anatomical differences were observed in LH198 or 3IIH6. B73 *rdla1* mutants had significantly fewer lateral veins per half leaf than wild-type (Table 1). While the trend for lower lateral vein counts was present in LH198, there was no significant difference in the number of lateral veins for LH198 or 3IIH6 (Table 1). Specific leaf area (SLA) was significantly higher in B73 and LH198 *rdla1* plants compared to wild-type, indicating that leaves in the mutant had less mass per unit area than wild-type (Table 1). SLA was not different between mutant and wild-type classes in the 3IIH6 genetic background. Neither percent carbon nor percent nitrogen in leaf tissue samples were significantly different between mutant and wild-type classes when measured in B73 (Table 1).

**Table 1:**
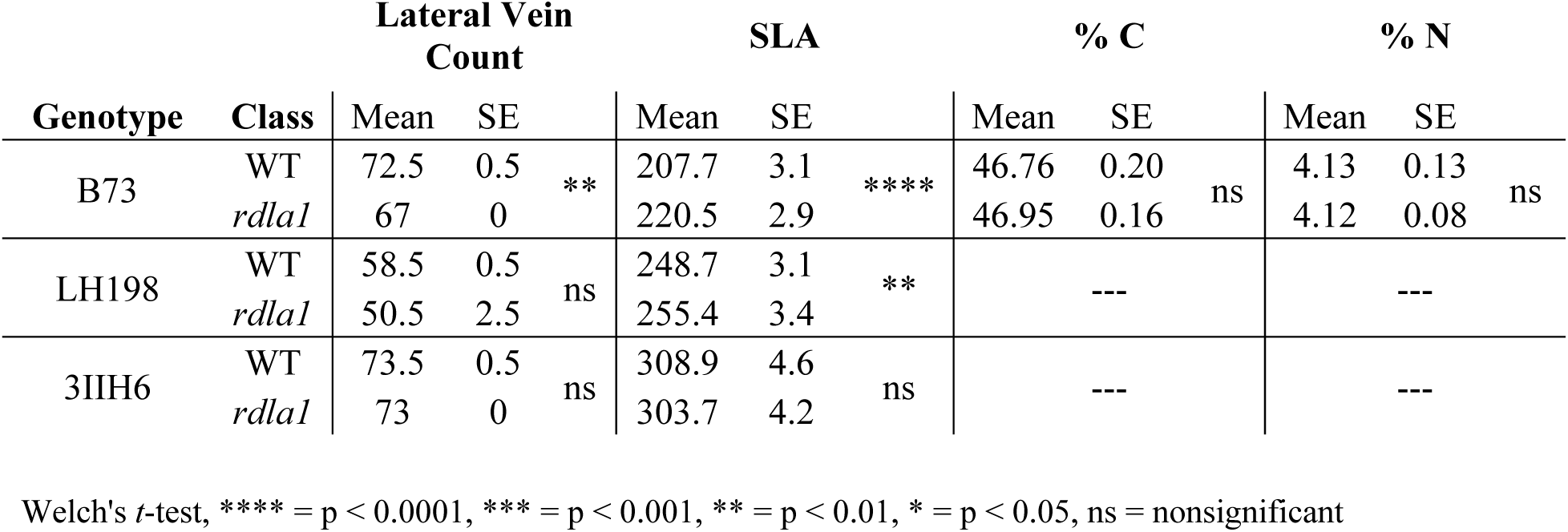
Leaf anatomical and physiological data collected on B73, LH198, and 3IIH6 wild-type and *rdla1* mutant introgression lines. Mean number of lateral veins on one side of the midrib of Leaf 2 ± standard error (n = 2 leaves). Mean specific leaf area (SLA) for Leaf 10 ± standard error (n = 4 blocks). Mean percent carbon for Leaf 10 ± standard error (n = 4 blocks). Mean percent nitrogen for Leaf 10 ± standard error (n = 4 blocks).

### Cuticular Wax Profiling

Because *Zm00001eb191580* is predicted to impact cuticular wax synthesis, we investigated the cuticular wax profile of B73 wild-type and *rdla1* plants. Of the 74 compounds detected by gas chromatography mass spectrometry (GC-MS) (Supplementary Table 3), heptacosane (C27 alkane) and nonacosane (C29 alkane) showed significantly elevated signatures for both juvenile (n = 6 pools, Welch’s *t*-test, p < 0.01) and adult (n = 6 pools, Welch’s *t*-test, p < 0.05) leaf tissue in *rdla1* mutants relative to wild-type (Figure 4a-b). Predicted precursor molecules of these alkanes are octacosanoic acid (C28 fatty acid) and triacontanoic acid (C30 fatty acid), respectively. No difference in octacosanoic acid was seen between *rdla1* and wild-type in juvenile leaf tissue, but a marginally insignificant reduction in this molecule was seen in adult leaves (n = 6 pools, Welch’s *t*-test, p > 0.05). Triacontanoic acid was not detected in juvenile leaves but was significantly enriched in adult wild-type leaves (n = 6 pools, Welch’s *t*-test, p < 0.05) (Figure 4c). A nonsignificant increase in hentriacontane (C31 alkane) was also detected for both juvenile and adult leaves in *rdla1* mutants, although the separation was more severe in adult leaves (n = 6 pools, Welch’s *t*-test, p > 0.05). Its precursor, dotriacontanoic acid (C32 fatty acid), was not detected at either growth stage. Three additional metabolites with differential accumulation patterns not clearly coupled to very long chain alkane biosynthesis were also detected. Tetracosanol (C24 primary fatty alcohol) and eicosanoic acid (C20 fatty acid) levels were significantly elevated in *rdla1* mutants relative to wild-type for adult and juvenile leaf stages, respectively (n = 6 pools, Welch’s *t*-test, p < 0.05) (Figure 4d-e). Beta-amyrin levels in adult leaves were significantly lower in *rdla1* mutants than wild-type (n = 6 pools, Welch’s *t*-test, p < 0.01) (Figure 4f).

**Figure 4:**
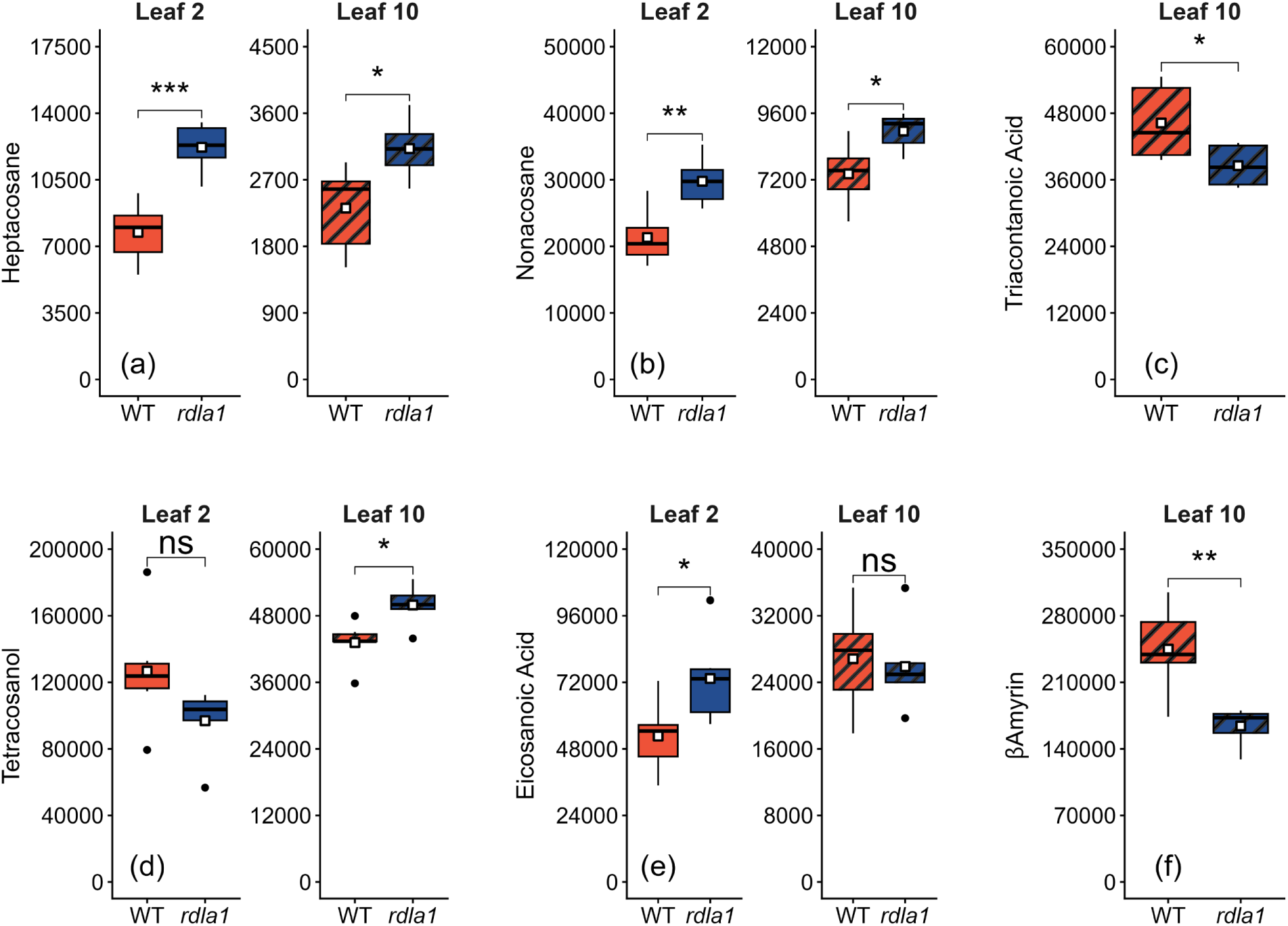
Boxplots of key metabolites collected from wild-type (orange) and mutant *rdla1* (blue) B73 plants. Samples were collected from juvenile (solid coloring) and adult (hatching) leaves. Metabolites reported are (a) heptacosane, (b) nonacosane, (c) triacontanoic acid, (d) tetracosanol, (e) eicosanoic acid, and (f) β-amyrin. The centerline of each boxplot indicates the median, while the white square indicates the mean. The box is bounded by the lower 25^th^ percentile and upper 75^th^ percentile. Whiskers extend to the largest and smallest values within 1.5× the interquartile range for the 75^th^ and 25^th^ percentiles, respectively, while points beyond whiskers represent values < 3× the interquartile ranges. Significant differences calculated using a two-tailed Welch’s *t*-test are denoted as follows: *** = p < 0.001, ** = p < 0.01, * = p < 0.05, ns = nonsignificant.

### Photosynthetic Properties

Since leaf morphology changes were observed in *rdla1* mutant lines, steady-state and *A/C_i_* curve gas exchange measurements were conducted to assess photosynthetic performance in B73, LH198, and 3IIH6 genetic backgrounds. No significant differences were observed for *A_net_, E, g_s_,* and *C_i_* between *rdla1* mutant and wild-type plants (n = 6 plants, Welch’s *t*-test, p > 0.05) (Supplementary Figure 5a-d). Furthermore, *iWUE*, calculated as *A_net_* divided by *g_s_*, was not significantly different between *rdla1* mutant and wild-type plants in any genetic background (n = 6 plants, Welch’s *t*-test, p > 0.05) (Supplementary Figure 5e). While *rdla1* plants trended as having lower *A_net_* than wild-type at each CO_2_ setpoint of the *A/C_i_* curve, there was no significant difference between mutant and wild-type plants for either *A_net_* or *C_i_* (n = 6 plants, Welch’s *t*-test, p > 0.05) (Supplementary Figure 5f). Collectively, photosynthetic metrics of *rdla1* and wild-type plants were highly similar when measuring traits on a leaf area basis.

**Figure 5:**
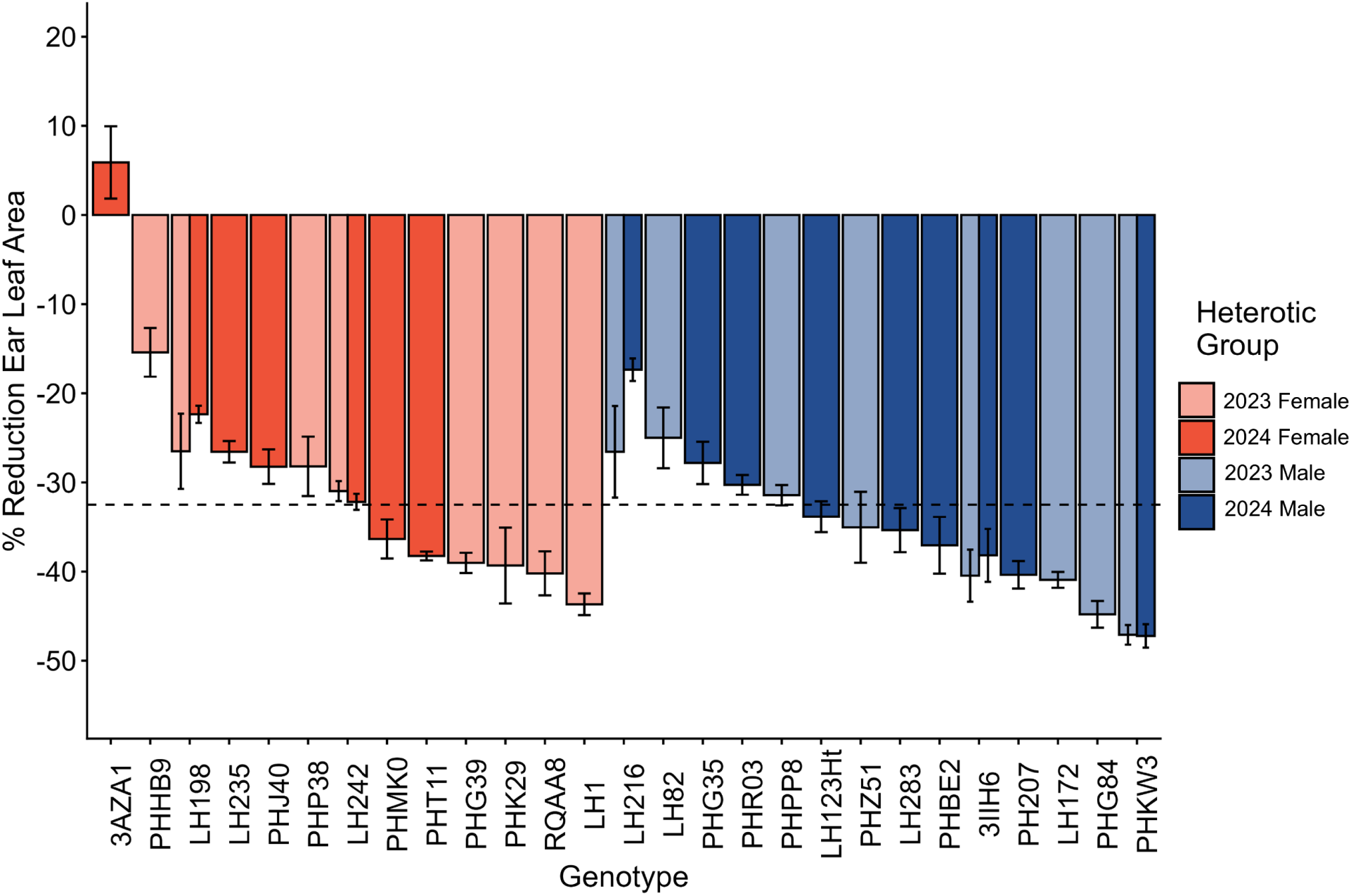
Average percentage leaf area reduction ± standard error of *rdla1* introgression lines relative to wild-type at the leaf of the ear-bearing node. Each bar represents the average of n = 4 plots (except 3IIH6 2023 is n = 2 plots). Orange bars are inbreds classified as female, and blue bars are inbreds classified as male. Genotypes with faded coloring were grown in the 2023 field trial, while genotypes with solid coloring were grown during the 2024 field trial. The dashed line represents the average percentage reduction in leaf area across all inbreds.

### ExPVP Panel Leaf

To evaluate the effect of the *rdla1* mutation across phenotypically diverse germplasm originating from commercial breeding programs, homozygous *rdla1* ExPVP introgression lines developed after five generations of backcrossing were grown in a split-plot design with homozygous wild-type segregant lines in 2023 and 2024. A total of 27 ExPVP backgrounds were evaluated. Sixteen ExPVP comparisons were planted in each field season, with five of the 27 total lines replicated in both years. While a significant year effect was observed for absolute ear leaf area values, there was no significant difference in the percentage reduction of leaf area across years within genotype (Supplementary Figures 6 & 7). Therefore, data from the two growing seasons were analyzed in aggregate. Across the ExPVP panel, changes in ear leaf area in *rdla1* introgressions relative to wild-type ranged from a slight increase of 5.90% ± 4.05% in 3AZA1 to a severe reduction of –47.23% ± 1.32% in PHKW3. The mean percentage reduction in leaf area across the panel was –32.51% (Figure 6). The range in absolute ear leaf area values for the wild-type class was 246cm^2^ ± 14 cm^2^ (3AZA1) to 647cm^2^ ± 29cm^2^ (PHK29), averaging 465cm^2^.The range in absolute ear leaf area values for the *rdla1* class was 196cm^2^ ± 11 cm^2^ (PHJ40) to 427cm^2^ ± 19 cm^2^ (PHHB9), averaging 309cm^2^.

**Figure 6:**
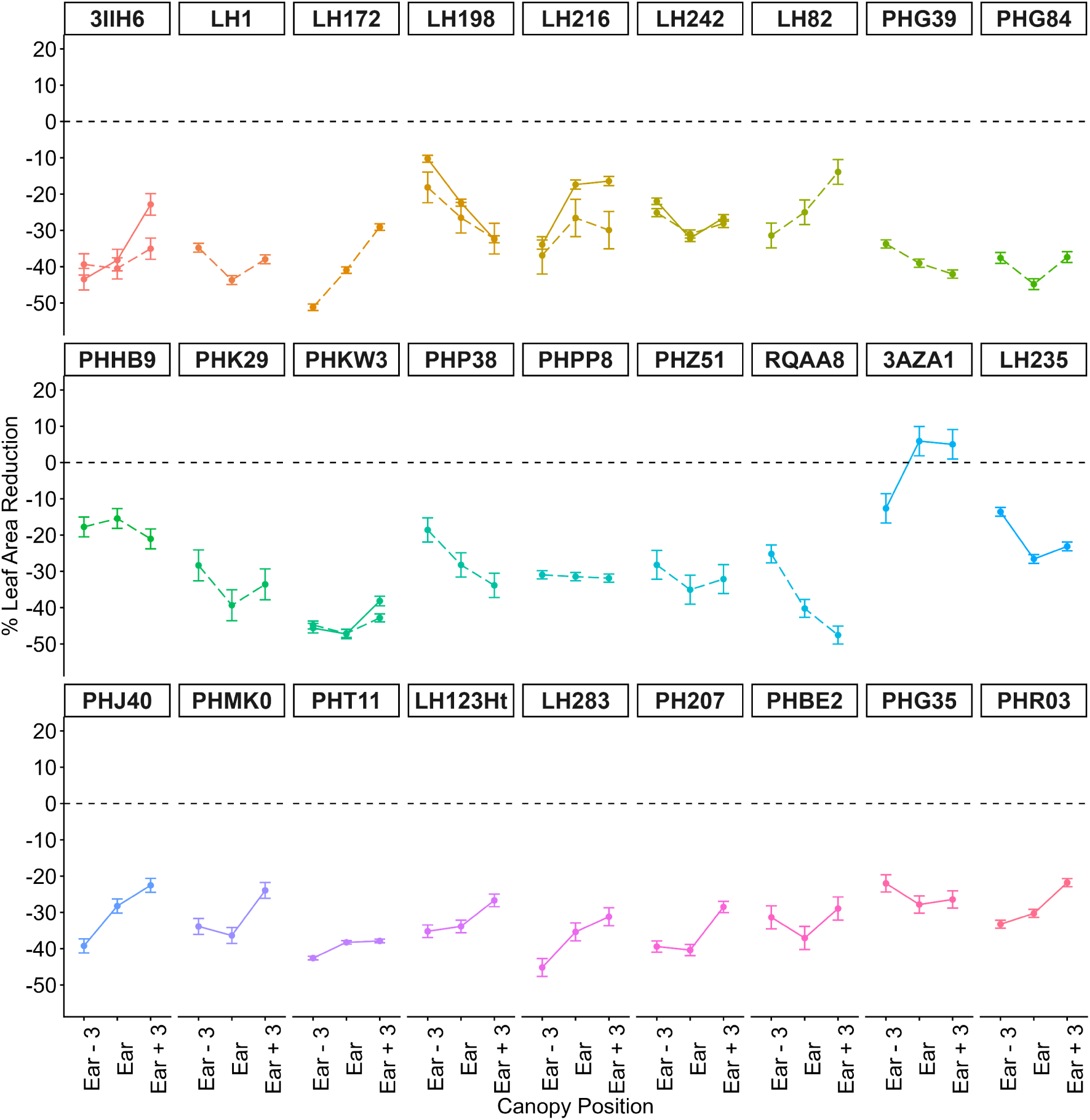
Average percentage leaf area reduction ± standard error of *rdla1* introgression lines relative to wild-type at the third leaf below the ear-bearing node, the leaf of the ear-bearing node, and the third leaf above the ear bearing node. Each point represents the average of n = 4 plots (except 3IIH6 2023 is n = 2 plots). Genotypes with dashed lines were grown in the 2023 field trial, while genotypes with solid lines were grown during the 2024 field trial.

The shape of a crop canopy is influenced by the size and dimensionality of all the plant’s leaves. In an attempt to practically quantify the distribution of leaf area effects within the canopy, leaves at the third node below and the third node above the ear-bearing node were compared to the ear leaf area in both genotypic classes (Supplementary Figure 8). As with B73, the effect size of ear leaf area reduction in *rdla1* lines was not directly proportional to the effect size at other positions within the canopy, with changes in rank order of ear leaf area reductions across the ExPVP panel at alternate canopy positions (Supplementary Figures 9a-b). For the third leaf below the ear-bearing node, percentage reduction in area ranged from –10.25% ± 0.96% in LH198 to –51.17% ± 0.89% in LH172 and averaged –31.41%. For the third leaf above the ear-bearing node, percentage change in area ranged from an increase of 5.01% ± 4.05% in 3AZA1 to a reduction of –47.54% ± 2.47% in RQAA8 and averaged –29.09%. Plotting the effect size of leaf area reduction in *rdla1* plants at these three canopy positions revealed distinct patterns in leaf area effect size that were conserved across years for replicated genotypes (Figure 6). In some lines, such as LH82 and LH172, the percentage reduction in leaf area was most extreme at lower canopy positions, with the effect size decreasing at higher leaf numbers. In contrast, lines like LH198 and PHP38 had greater reductions in leaf area in upper canopy positions. For PHPP8, the percentage reduction in leaf area was stable regardless of canopy position. Thus, a single mutation gives rise to a spectrum of plant architecture changes that can influence the dynamics of canopy shape and light interception.

## Discussion

Through fine-mapping and sequencing-based approaches, we demonstrated that an insertion allele of *Zm00001eb191580* underlies the *rdla1* phenotype. *Zm00001eb191580* bears closest homology to *ECERIFERUM1* (*CER1*), where it shares 53.27% protein identity with *Zm00001eb247450* (*ZmCER1*) and 52.83% protein identity with *At1g02205* (*AtCER1*). The *ECERIFERUM* (*CER*) gene family is highly divergent in *Arabidopsis thaliana*, with 20 functionally characterized *CER* genes^31,32^. Visible phenotypes of these mutants often include a shiny appearance of the leaves, and these genes have previously been identified as analogous to some members of the *GLOSSY* (*GL*) gene family in maize^33,34^.

In the leaf cuticular wax biosynthetic pathway of *Arabidopsis*, precursor very long chain fatty acids are reduced to very long chain aldehydes by a suite of ECERIFRUM proteins including CER3, CER8, and CER9. These aldehydes may either be diverted into the production of primary alcohols via CER4, which can subsequently be used to produce high molecular weight esters, or very long chain fatty alkanes via CER1 or CER16, with a fraction of alkanes then used to make secondary alcohols and ketones^31^. In *rdla1* plants, C27 and C29 alkanes were significantly enriched, while the precursor molecules of these alkanes, C28 and C30 fatty acids, were depleted. These results suggest that there is a conservation of CER1 function between maize and *Arabidopsis*. Under this model, *Zm00001eb191580* and *Zm00001eb247450* compete for similar precursor very long chain fatty acid molecules, and loss of function of *Zm00001eb191580* in *rdla1* mutants diverts more of these substrates to CER1. However, no metabolites were identified through GC-MS that were likely products of RDLA1, whether alkanes or downstream molecules. As such, the current functional activity and gene products of RDLA1 remain unknown. Regulating cuticular wax biosynthesis represents a novel pathway for manipulating leaf area in maize, yet has strong foundations from observed interactions of wax profiles and reduced leaf area in both *Arabidopsis* and rice^35,36^.

Existing studies have functionally investigated few of the *CER* orthologs in maize^34^. Knockout mutants of *ZmCER1* have been characterized as producing a similar phenotype to that of *rdla1* mutants, both in visual appearance and quantitative measurements of leaf area. Maize *cer1* mutants produced leaves with reductions of length and width at the ear-bearing node that aligned closely with our observations of *rdla1*, and a crinkled appearance that was attributed to aberrant bulliform cell development^37^. Contrary to these reports for *cer1* mutants, we did not find the leaves of *rdla1* plants to have abnormal bulliform cell arrangement in transverse sections of leaf tissue. Leaves were developmentally complete and displayed normal morphology, apart from having fewer lateral veins than wild-type. It is possible that the lack of an obvious cellular phenotype was due to sampling stage. While *rdla1* mutants show a quantitative reduction in leaf area even as early as Leaf 2, the visual crinkled leaf phenotype characteristic of this mutation does not arise until after the transition of juvenile to adult leaf tissue at approximately Leaf 6 in B73.

In multiple genotypes, we found elevated SLA values in the *rdla1* mutants, suggesting a deficit in cuticular wax accumulation that would contribute to reduced leaf mass. Consistent with this hypothesis, knockout and overexpression lines of *ZmCER1* had decreased and increased wax deposition, respectively^37^. However, no analysis was completed to determine any alterations to the mutant wax profile in the *cer1* mutants evaluated in that study. Interestingly, *cer1/rdla1* double mutants showed characteristic leaf crinkling immediately after emergence and did not survive past the Leaf 3 growth stage^37^. These results indicate that *ZmCER1 and RDLA1* share functional overlap in biological processes essential for leaf development.

Owing to the highly aliphatic nature of the wax layer, the primary function of epicuticular wax is to protect leaves from desiccation. As such, cuticular wax mutants have often been associated with increased water loss and susceptibility to other abiotic and biotic stressors^37,38,39^. We noted no significant differences in leaf water loss via transpiration during steady-state gas exchange measurements. However, the relative contribution of cuticular transpiration to total plant water loss is orders of magnitude less than the primary driver of plant water loss, stomatal conductance^40^. Thus, any impact of *rdla1* on cuticular permeability was likely not detectable by our standard gas exchange approach and remains open to investigation. Nevertheless, from an agronomic water use perspective, we may assert that differences in water loss affecting normal plant growth and development are negligible.

We noted minimal pleiotropy for the physiological and agronomic traits monitored in this study, including total plant height, height of the ear-bearing node, %C and %N in leaf samples, and photosynthetic performance. In B73, we found that the greatest effect size of the *rdla1* mutation was in the middle of the canopy near the ear-bearing node, where the absolute area of leaves is greatest. Leaves at the ear-bearing node consistently had the largest absolute area across all genotypes evaluated, yet the effect size of the mutation was not always greatest at this position. Notably, genotypes with the largest absolute leaf areas did not always show the largest leaf area reduction with the *rdla1* mutant allele. The relative effect of the mutation within genotype across years was conserved, indicating that the *rdla1* mutant allele shows stable performance across environments.

Previous studies have identified leaf area traits as key effectors of plant productivity, driving the discovery of associated QTLs for these traits^18,41,42,43^. In an optimized crop canopy, leaves near the top of the plant should not shade the middle and lower canopy positions, allowing leaves throughout the plant to contribute to photosynthetic productivity^16,20^. Genotypes like LH198, PHP38, and RQAA8, which show more severe leaf area reductions in the upper canopy when the *rdla1* allele has been introgressed, should act to minimize this shading and allow deeper light penetration to lower positions. Genotypes including LH172, PHJ40, and LH283 instead show significantly more severe leaf area reductions at lower canopy positions, indicating that these genotypes may show more limited utility at higher planting densities. Nevertheless, the existence of differential, even inverse, response patterns to *rdla1* across broad genetic backgrounds demonstrates the potential for using this allele as a tool for selecting idealized canopy structure within maize breeding programs. Indeed, the differences in expressivity of the *rdla1* allele offer a unique opportunity to characterize a novel genetic regulatory network of loci interfacing leaf-level developmental physiology with canopy-level architectural traits. Such findings will be invaluable for designing an ideotype canopy structure optimized throughout developmental time. We posit that, due to its moderate effect on leaf area across genetic backgrounds with limited pleiotropic effects, the *rdla1* allele may serve as a future target for optimization of leaf area for high-density cropping systems. Because maize is grown primarily as a hybrid crop, claims about novel phenotypes impacting plant productivity and grain yield should be made in the context of hybrids rather than inbreds alone^44^. The development of *rdla1* hybrids from the genotypes included in our ExPVP introgression panel, which represent a subset of the most elite germplasm available for public-sector research programs, will play a key role in assessing the impact of this mutation on density tolerance and grain yield in an agronomically relevant setting.

## Methods

### DNA Extraction, Sequencing, and Variant Analysis

DNA was extracted using a modified CTAB protocol from frozen or lyophilized seedling or adult leaf tissue^45^. Whole-genome shotgun library and sequencing was done by the at the Roy J. Carver Biotechnology Center at the University of Illinois using the Hyper Library construction kit (Kapa Biosystems, Wilmington, MA). For two whole-genome sequenced (WGS) libraries, DNA from four Oh43 WT and four *rdla1* BC_5_S_10_ plants was pooled separately and sequenced on an Illumina HiSeq4000 (Illumina, San Diego, CA) lane producing paired reads of 150 bp. For the third WGS library, DNA from 17 BC_6_S_1_ *rdla1* plants was normalized and bulked, then sequenced on an Illumina NovaSeq6000 SP (Illumina, San Diego, CA) lane producing single reads of 100 bp. The 17 BC_6_S_1_ *rdla1* plants were selected for informative recombination events in the *rdla1* region using sequencing or PCR markers from the 208 BC_6_S_1_ *rdla1* individuals genotyped in the study.

Restriction site associated DNA sequencing (RAD-seq) libraries were prepared as described^46^. The referenced protocol is an adaptation from an earlier established protocol^47^. For the first set of RAD-seq libraries, DNA from four Oh43 WT plants and 26 BC_6_S_1_ *rdla1* plants was digested separately using two enzyme combinations, *Ape*KI*-Pst*I and *Hin*P1I-*Pst*I. Single reads of 150 bp were obtained for the RAD-seq libraries, which were sequenced on an Illumina HiSeq4000. RAD-seq libraries were also built for the 17 recombinant bulk whole genome-sequenced BC_6_S_1_ *rdla1* individuals described above using three enzyme sets, *Ape*KI*-Pst*I, *Hin*P1I*-Pst*I, and *Ape*KI only. The libraries were sequenced on an Illumina NovaSeq6000 SP lane for single reads of 100 bp.

FASTQ files were demultiplexed with the *process_radtags* command in Stacks 2.41 if appropriate^48^. Reads were aligned to the maize Zm-B73-REFERENCE-GRAMENE-4.0 (B73) reference genome and also to the maize Zm-Oh43-REFERENCE-NAM-1.0 (Oh43) reference genome using Bowtie 2.3.2^49^. Coordinates were subsequently converted to Zm-B73-REFERENCE-NAM-5.0. Genotype probability calculations and variant calls were performed in BCFtools 1.7^50^. Variant filtering was conducted in BCFtools 1.7 and VCFtools 0.1.15^50,51^. Throughout, only the two most common alleles per site were considered.

Variant filtering varied by analysis. To map the *rdla1* region to a chromosome, histograms of the physical positions of sites which were polymorphic between the whole-genome sequenced WT and BC_5_S_10_ *rdla1* were constructed. The B73 alignment was used, and only homozygous sites with average depth greater than ten present in both individuals were used. To subsequently fine-map *rdla1* within the region identified on chromosome four in the BC_6_S_1_ generation, all sequencing information from all individuals was used. For fine-mapping, the Oh43 alignment was used because a gap between contigs was present in the *rdla1* region in the B73 reference genome. Sites were classified as *rdla1*-associated if at least one call each of the WT and *rdla1* genotype was available, the site segregated among WT and *rdla1* individuals but was fixed within WT and within *rdla1* individuals, and the site was homozygous for the non-reference allele within the *rdla1* individuals. Histograms of the physical positions of *rdla1*-associated sites were generated in R. Variant effect prediction was conducted in the SnpEff 4.3 eff plugin on Galaxy for sites in the 151 kb region on Chromosome four to which *rdla1* ultimately mapped^52,53^.

### PacBio Long-Read Sequencing

High molecular weight genomic DNA was isolated by the Roy J. Carver Biotechnology Center at the University of Illinois from nuclei extracted from coleoptiles of a Oh43 *rdla1* BC_5_S_11_ seed stock. The DNA was sheared with a Megaruptor 3 (Diagenode Inc., Denville, NJ) to an average fragment length of 13kb, which were used to prepare a library with the SMRTBell Prep kit 3.0 (PacBio, Menlo Park, CA). The library was sequenced on 2 SMRTcell 8M on a PacBio Sequel IIe (PacBio, Menlo Park, CA) using the CCS sequencing mode and a 30hs movie time. These produced nearly three million reads with a mean read length of 14.4kb. CCS analysis was done in instrument with SMRTLink V11.1 using the following parameters:ccs --min-passes 3 --min-rq 0.99. Custom scripts were then used to isolate reads that encompassed the five gene models in the final fine mapping region. These reads were assembled into a single scaffold using hifiasm^54^. The resulting scaffold was compared to the reference genome for large structural differences.

### Plant Materials & Marker-Assisted Backcrossing

The *rdla1* mutation was originally isolated from a heterogeneous population and stabilized by backcrossing five generations with phenotypic selection into an Oh43 inbred background and then selfing for ten generations. This Oh43 *rdla1* conversion was used as a donor stock for introgression into B73 and a panel of 27 Expired Plant Variety Protection (ExPVP) inbreds (Supplementary Table 2) obtained from the USDA Germplasm Resources Information Network (GRIN). Marker-assisted selection was used to advance individuals carrying the *rdla1* haplotype during the backcrossing process. After five generations of backcrossing (BC_5_), individuals heterozygous for markers within the *rdla1* region were self-pollinated to isolate wild-type and *rdla1* progeny for the development of homozygous stocks. All stocks used for phenotyping experiments were advanced to the BC_5_S_2_ or BC_5_S_3_ stage.

Markers were designed using Primer3 through NCBI Primer Blast^55^ (Supplementary Table 1). Most primers were designed flanking the introns of annotated genes within the candidate interval to identify polymorphisms. PCR products were amplified via GoTaq Flexi (Promega, Fitchburg, WI) polymerase according to the following reaction conditions: 95°C for 2 minutes, 34 cycles of 94°C for 30 seconds, 59°C for 30 seconds, 72°C for 30 seconds, and 72°C for 10 minutes. PCR products were scored after running on agarose gels ranging 1.2% to 4% depending on fragment length.

### Inbred Characterization

In the 2023 field season, 18 pairs of inbreds were grown, and in the 2024 field season, 16 pairs of inbreds were grown (Supplementary Table 2). Both experiments were located at the University of Illinois Vegetable Crops Research Farm in Champaign, IL. Inbreds were grown in a split-plot design, with single rows of WT and *rdla1* versions of the same inbred grown adjacent to each other. Row spacing was 0.76 m with 3.66 m plot length and a target planting density of 11,736 plants ha^−1^. Each genotype was replicated across four blocks unless otherwise noted. For B73, the length and width of each leaf from stages V2 to V10 was measured to the nearest millimeter with a ruler. Area was calculated as the length multiplied by the width multiplied by a conversion factor of 0.75. The length, width, and area of each leaf from stages V9 to V20 was measured using a LI-3000C (LI-COR Biosciences, Lincoln, NE) leaf area meter. For all other inbreds, leaf length, width, and area measurements were collected on the leaf of the ear-bearing node, the third leaf above the ear-bearing node, and the third leaf below the ear-bearing node with the leaf area meter. Ten plants from each row were measured as technical replicates for leaf area. Plant height and ear height were measured on five plants per plot post-flowering as technical replicates using a plant height stick marked in 1cm intervals.

### Leaf Transverse Sectional Imaging

Seeds of B73, 3IIH6, and LH198 wild-type and *rdla1* mutant introgression lines were grown in greenhouse conditions at Cornell University until the V2 growth stage. Leaf 2 was harvested from two individuals of each genotype, fixed in 3.7% formalin/acetic acid/alcohol (FAA), and paraplast embedded as described^56^. Images of 10μm transverse histological sections stained with safranin-fast green staining were obtained using established protocols with adaptations as previously described^57^. This protocol was an adapted from Micrograph images were obtained on a Zeiss Z1-Apotome microscope (Zeiss, Oberkochen, Baden-Württemberg, Germany). Measurements were made using Zeiss Axiovision software release 4.6. The number of lateral veins was counted manually on one side of the midrib of all leaves, and morphology of the cells was assessed visually.

### Leaf Metabolites

In the 2025 field season, B73 wild-type and *rdla1* mutant plants were grown in a single replicate of five-row plots at the University of Illinois Vegetable Crops Research Farm in Champaign, IL. Row spacing was 0.76 m with 3.66 m plot length and a target planting density of 11,736 plants ha^−1^. At the V2 growth stage, the entire second leaf was harvested from five individual plants to form a pooled biological replicate with enough tissue for analysis. A total of six biological replicates were harvested per genotype. At the V10 growth stage, a circular cork borer with diameter of 1.77 cm was used to collect four leaf discs from the tenth leaf of five individual plants to form a pooled biological replicate. Once again, six biological replicates were harvested per genotype. Immediately after harvest, tissue was flash frozen in liquid nitrogen. Samples were shipped to the University of Missouri Metabolomics Center (Columbia, MO) for cuticular wax extraction and quantification. Cuticular waxes were extracted in chloroform and analyzed by gas chromatography – mass spectrometry (GC-MS) based on an adapted protocol previously described^59^.

### Gas Exchange Measurements

At ∼V10 growth stage, gas exchange was conducted on field-grown plants using a LI-6800 (LI-COR Biosciences, Lincoln, NE) gas exchange analyzer. The cuvette was fixed with a 6 cm^2^ clamp and flow rate set to 500 μmol s^−1^. Light intensity was 2000 μmol mol^−1^ with a red / blue ratio of 10% / 90%. Cuvette conditions were controlled with relative humidity of 60%, CO_2_ concentration of 400 μmol mol^−1^, and leaf temperature of 28°C. All measurements were collected on the uppermost fully expanded leaf. After clamping, leaves were acclimated to cuvette conditions for ∼30 minutes to reach steady-state. For B73, LH198, and 3IIH6, steady-state data were logged every 10 seconds for five minutes and averaged. A total of n = 6 biological replicates were measured per reported genotype × class combination. Additionally for B73, *A/C_i_* curves were constructed following 15 CO_2_ concentration points: 400, 300, 200, 100, 50, 10, 25, 75, 150, 250, 400, 600, 800, 1000, and 1200 μmol mol^−1^.

### Specific Leaf Area, %C, and %N

At ∼V10 growth stage, a circular cork borer with diameter of 1.77 cm was used to collect leaf discs from the uppermost fully expanded leaf from WT and *rdla1* B73, 3IIH6, and LH198. Four technical replicates were collected from each plant, and ten plants were sampled per plot as biological replicates. Leaf punches were dried at 65°C for seven days to achieve shelf stability; immediately prior to measurement, leaf punches were again dried at 65°C for seven days. Leaf punches were weighed to the nearest 0.0001 g on a NewClassic MF (Mettler Toledo, Zurich, Switzerland) analytical balance. Specific leaf area (SLA) was calculated as the area of the disc divided by the dry mass. After recording specific leaf area, leaf discs from B73 wild-type and *rdla1* plants were prepared for carbon and nitrogen analysis. Dried leaf tissue from three plants per plot were pooled together into a single tube and ground using a 1600 MiniG grinder (SPEX SamplePrep, Inc., Metuchen, NJ), generating three technical replicates per plot. Samples were analyzed for %C and %N via mass spectrometry at the Washington State University Stable Isotope Core Laboratory (Pullman, WA).

### Complementation Test

UFMu-09400 was obtained from the Maize Genetics Cooperation Stock Center (Urbana, IL) as a stock carrying *mu1077278* in *Zm00001eb191580*. During the summer of 2024 field season, a plant heterozygous for *mu1077278* was crossed with a homozygous PHKW3 *rdla1* plant. Kernels from this cross were planted in a 50-cell tray at the University of Illinois Plant Care Facility greenhouses. Following genotyping for both *mu* and *rdla1* alleles, eight individuals of each genotype *rdla1/mu* and *rdla1/+* were transplanted into a ground bed two weeks after germination. Plants were grown to maturity under supplemental LED lighting set to 12-h day length and 29/23°C maximum day/night temperatures. After transplanting, plants were fertilized with ammonium sulfate (21-0-0-24) and with supplemental foliar application of Sprint 330 chelated iron. At flowering, leaf length, width, and area measurements were taken on the leaf of the ear-bearing node, the third leaf below the ear-bearing node, and the third leaf above the ear-bearing node using a LI-3000C (LI-COR Biosciences, Lincoln, NE) leaf area meter.

### Statistical Analyses

All statistical analyses were conducted in R (version 4.5.2) using RStudio^60,61^. All comparisons were paired between wild-type and *rdla1* mutant classes of the same inbred genotypic background. When assumptions of normality and equivalence of variance for the data were met, a two-tailed Welch’s *t*-test was implemented using the ‘t.test()’ function.

## Acknowledgements and Funding

We thank Dr. Robert Lambert for providing the original mutant stock containing the *rdla1* allele. We also thank Mark A. Mikel for guidance on selection of the germplasm to be used for the ExPVP introgression panel. Furthermore, we thank the University of Illinois Roy J. Carver Biotechnology Center for conducting the PacBio sequencing for structural variant identification, and Dr. Allison Kolbe for assistance with long-read sequence analysis. Special thanks is given to the University of Missouri Metabolomics Center for performing the cuticular wax extractions and to the Washington State University Stable Isotope Core Laboratory for analyzing %C and %N. This research was performed with support from the Center for Research on Programmable plant Systems (CROPPS) under National Science Foundation Grant No. DBI-2019674.

## Author Contributions

M.J.R., M.R.L., and A.J.S. conceived the project and developed the associated germplasm. M.J.R. and A.J.S designed the field experiments. M.R.L. and M.J.R mapped the causal allele. M.J.R. performed the complementation test. M.J.S. performed the transverse microscopy. M.J.R. collected data for gas exchange, SLA, and leaf metabolites. M.J.R. and M.I.A. measured leaf area for the inbred introgression panel. M.J.R. analyzed the data. M.J.R. and A.J.S. wrote the manuscript. All authors reviewed and revised the manuscript.

## Abbreviations

*A_net_*: net CO_2_ assimilation
*CER*: ECERIFERUM
*C_i,_* intercellular: CO_2_ concentration, *E,* transpiration
ExPVP: Expired Plant Variety Protection
*g_s_*: stomatal conductance to water
*iWUE*: intrinsic water-use efficiency
rdla1: reduced leaf area1
SLA: specific leaf area

## Supplementary Figures

**Supplementary Figure 1:**
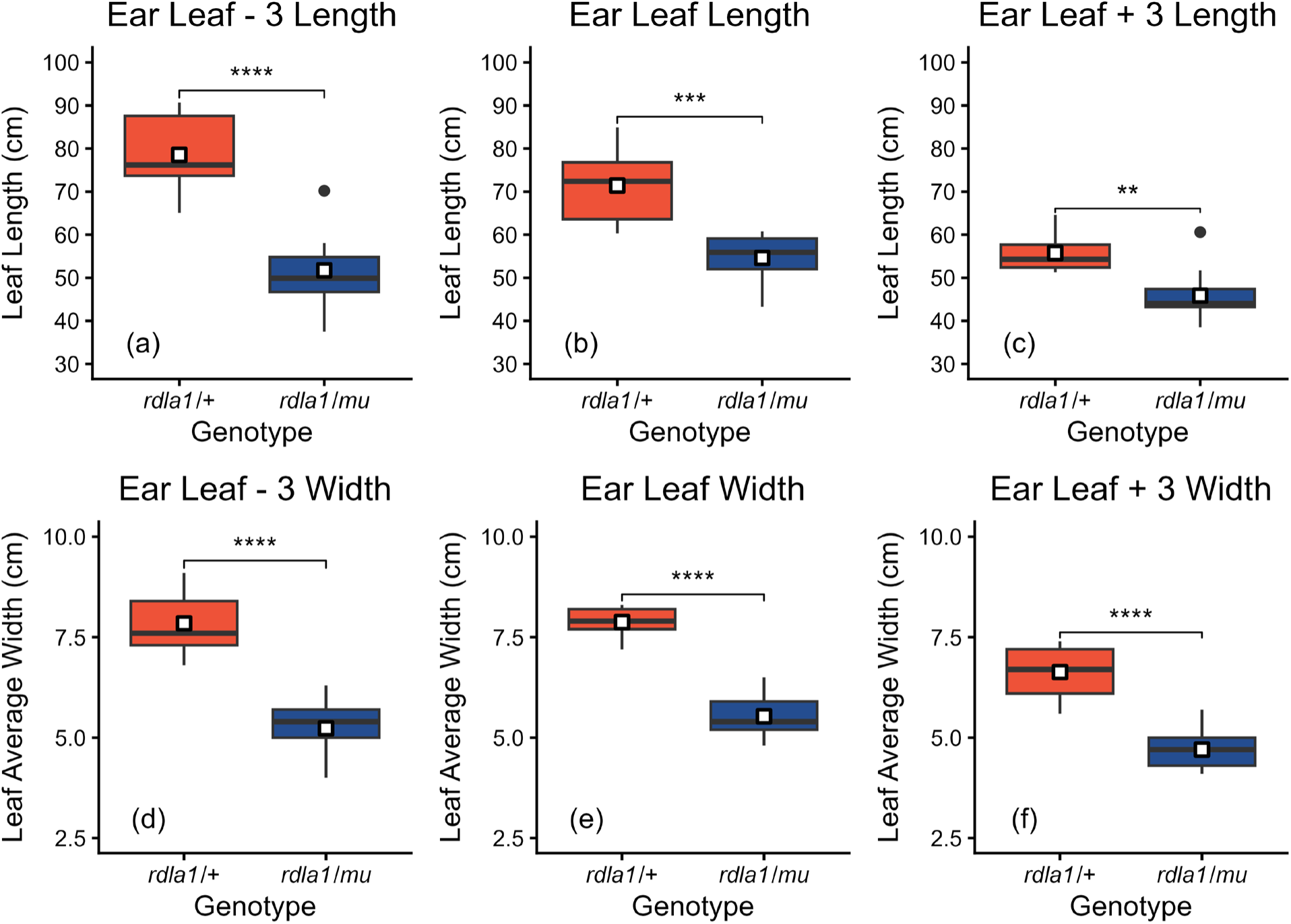
Absolute leaf length (a – c) and leaf width (d – f) reduction of *rdla1/+* (orange) and *rdla1/mu1077278* (blue) plants at the third leaf below the ear-bearing node (a & d), the leaf of the ear-bearing node (b & e), and the third leaf above the ear bearing node (c & f) (n = 8 plants). The centerline indicates the median, while the white square indicates the mean. The box is bounded by the lower 25^th^ percentile and upper 75^th^ percentile. Whiskers extend to the largest and smallest values within 1.5× the interquartile range for the 75^th^ and 25^th^ percentiles, respectively, while points beyond whiskers represent values < 3× the interquartile ranges. Significant differences calculated using a two-tailed Welch’s *t*-test are denoted as follows: **** = p < 0.0001, *** = p < 0.001, ** = p < 0.01, * = p < 0.05, ns = nonsignificant.

**Supplementary Figure 2:**
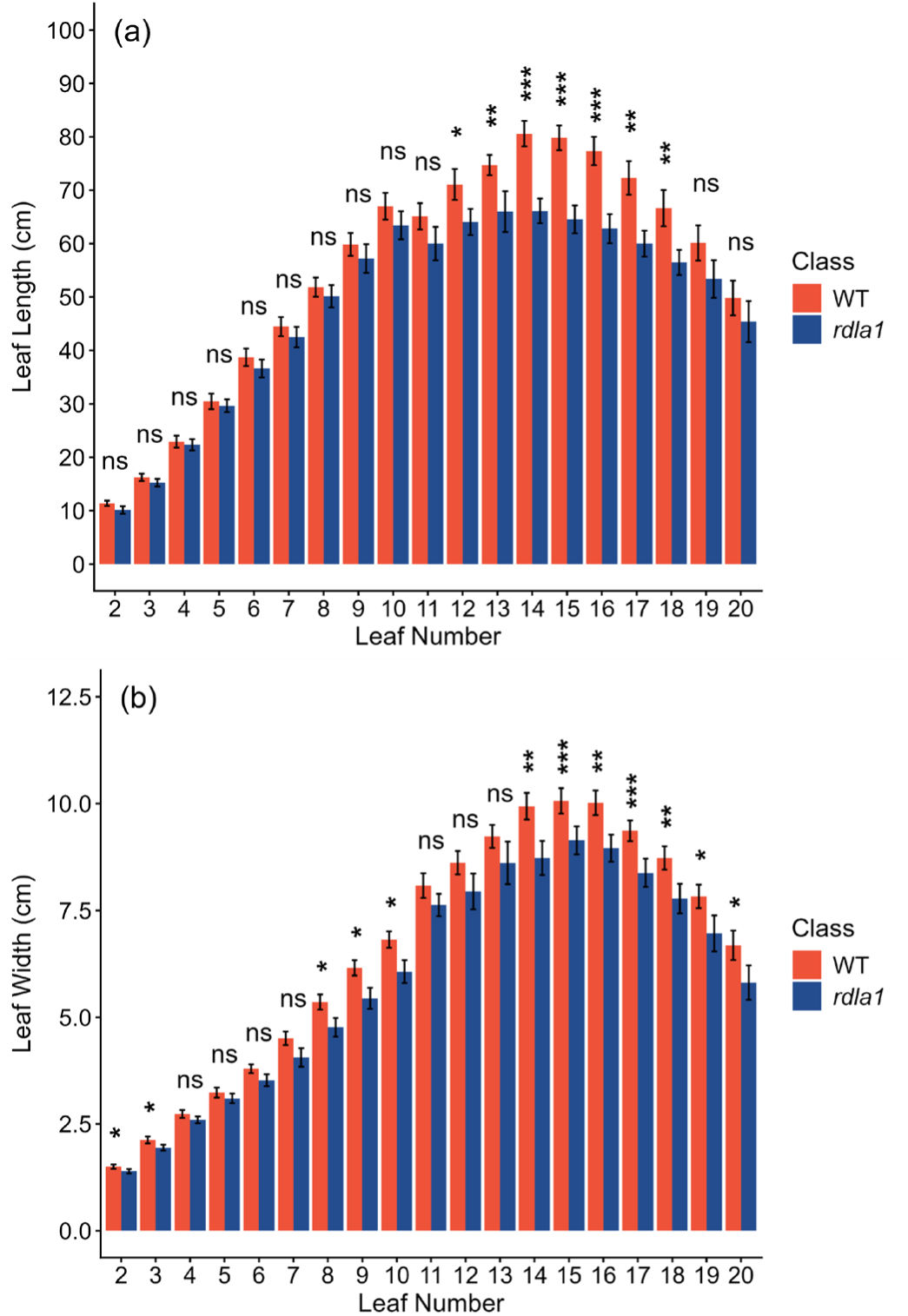
The mean leaf length (a) and mean leaf width (b) of wild-type (orange) and *rdla1* (blue) leaves for leaf positions of Leaf 2 through Leaf 20. Bars denoting mean leaf length and width are representative of plots (n = 4) ± standard error. Significant differences calculated using a two-tailed Welch’s *t*-test are denoted as follows: *** = p < 0.001, ** = p < 0.01, * = p < 0.05, ns = nonsignificant.

**Supplementary Figure 3:**
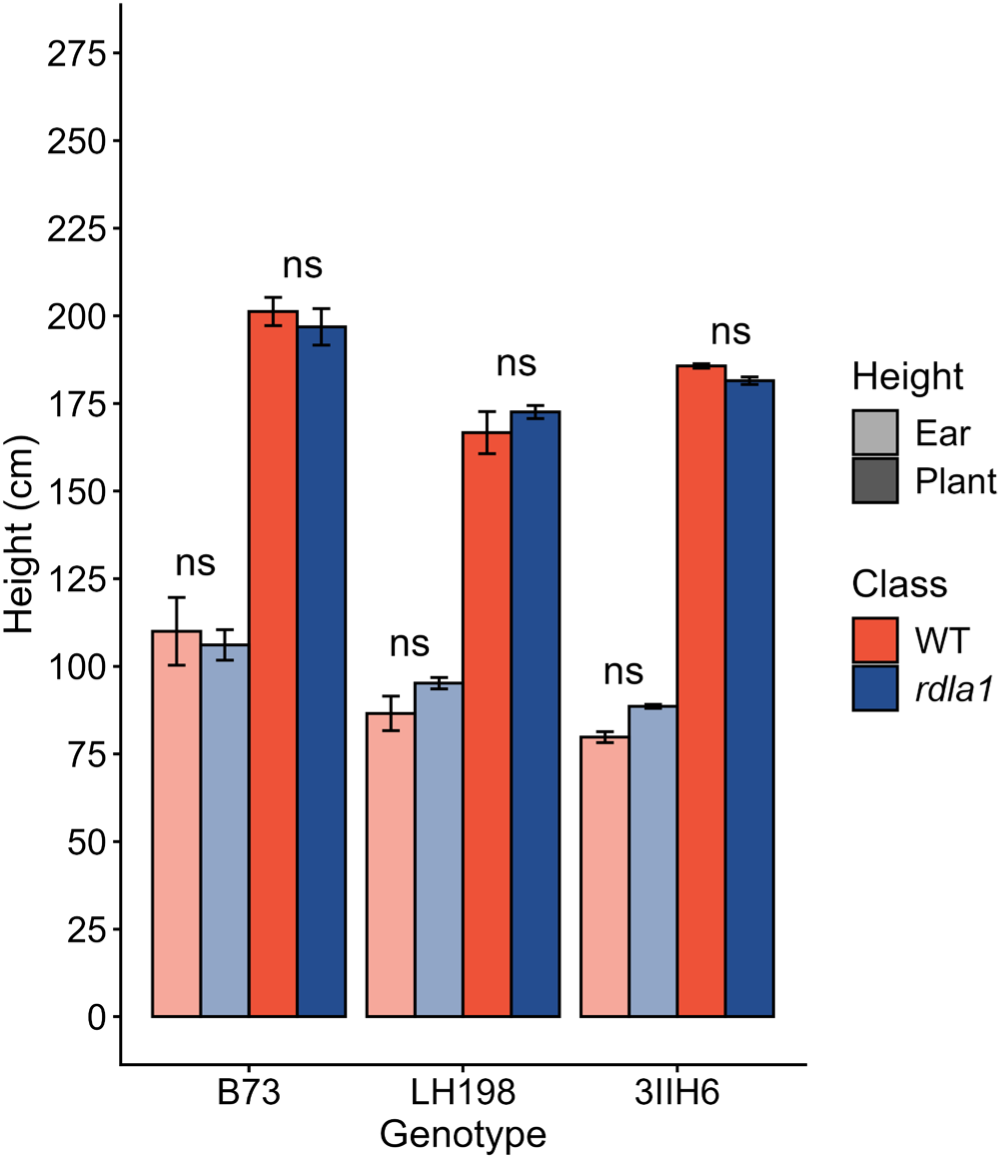
The mean height of the ear-bearing node (faded) and mean total plant height (solid) of wild-type (orange) and *rdla1* (blue) plants. Bars denoting mean height measurements are representative of plots (n = 4 for B73 and LH198, n = 2 for 3IIH6) ± standard error. Significant differences calculated using a two-tailed Welch’s *t*-test are denoted as follows: *** = p < 0.001, ** = p < 0.01, * = p < 0.05, ns = nonsignificant.

**Supplementary Figure 4:**
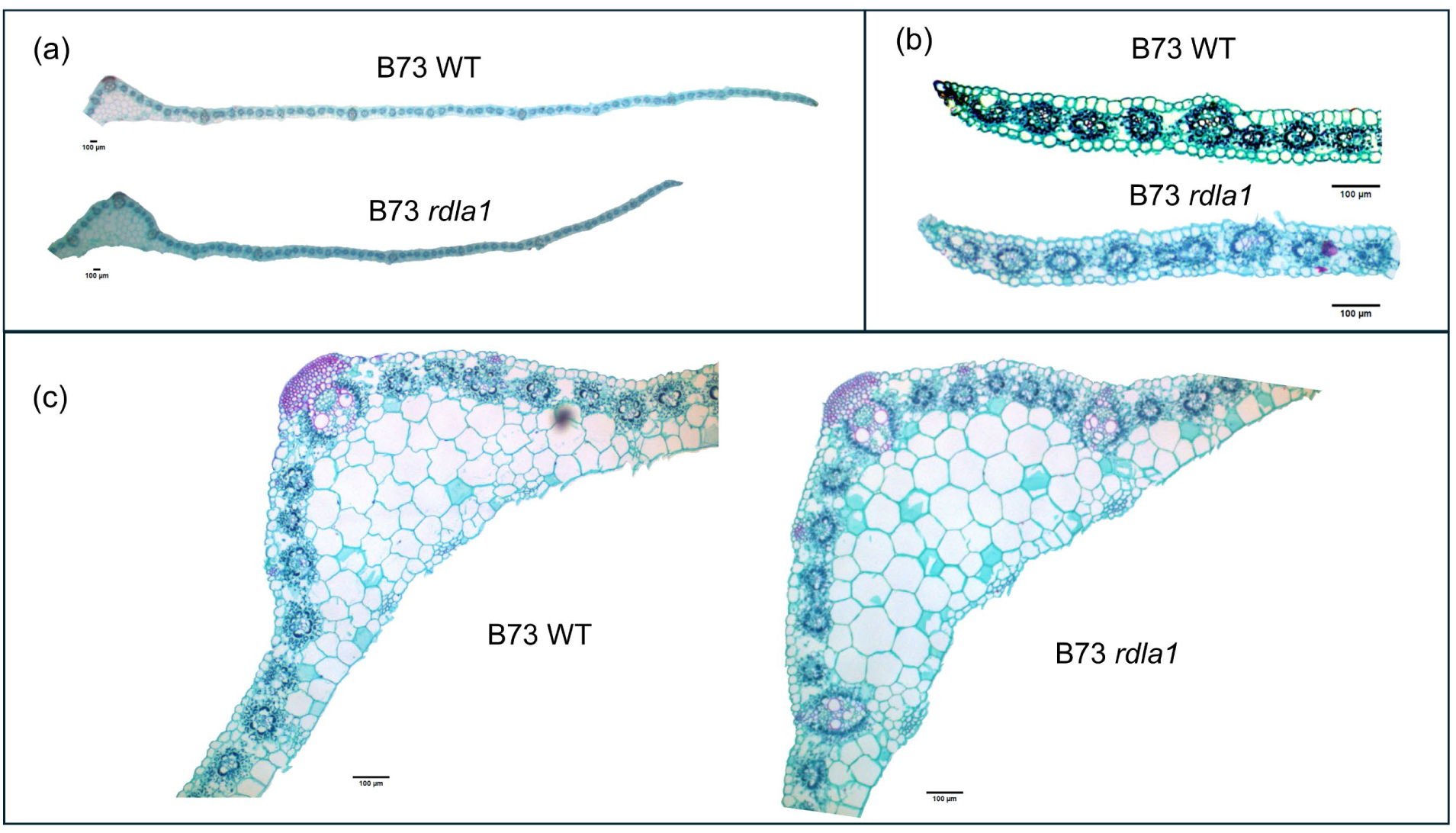
Histological imaging of B73 wild-type and *rdla1* mutant seedling leaves. (a) Seedling half-leaves reveal that *rdla1* mutant seedlings are slightly narrower than wild-type. Closer images of leaf margins (b) and midribs (c) reveal normal morphology and anatomy in *rdla1* mutant leaves. Scale bars = 100μm.

**Supplementary Figure 5:**
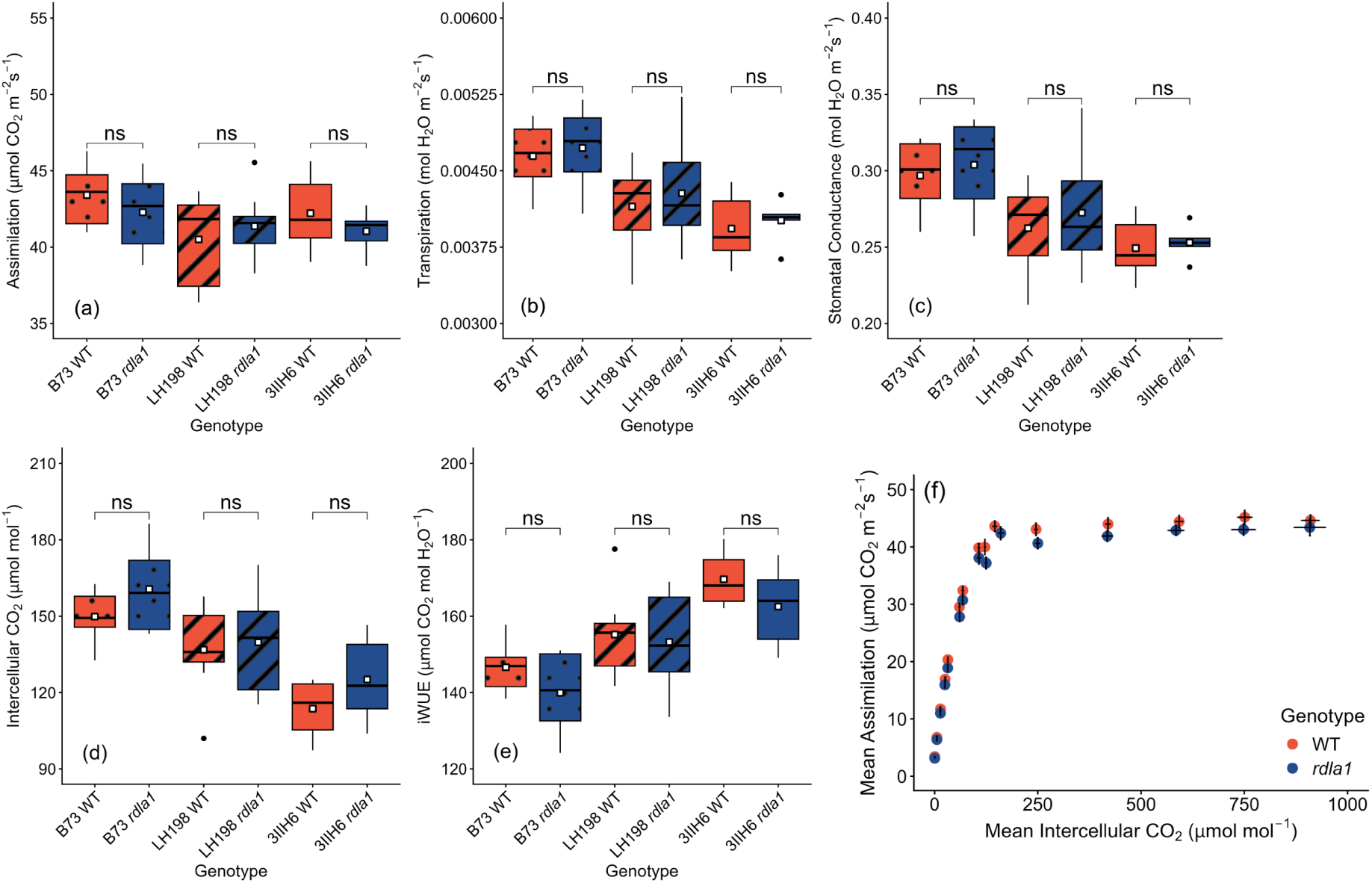
Boxplots of steady-state gas exchange (a-e) of B73, LH198, and 3IIH6 wild-type (orange) and *rdla1* (blue) plants. (a) Net CO_2_ assimilation; (b) transpiration; (c) stomatal conductance to water; (d) intercellular CO_2_ concentration; (e) intrinsic water-use efficiency. The centerline of each boxplot indicates the median, while the white square indicates the mean. The box is bounded by the lower 25^th^ percentile and upper 75^th^ percentile. Whiskers extend to the largest and smallest values within 1.5× the interquartile range for the 75^th^ and 25^th^ percentiles, respectively, while points beyond whiskers represent values < 3× the interquartile ranges. An *A/C_i_* curve (f) was also measured for B73 wild-type (orange) and *rdla1* mutant (blue) plants. Points on the *A/C_i_* curve are shown ± standard error for both *A* and *C_i_*. Significant differences calculated using a two-tailed Welch’s *t*-test are denoted as follows: *** = p < 0.001, ** = p < 0.01, * = p < 0.05, ns = nonsignificant.

**Supplementary Figure 6:**
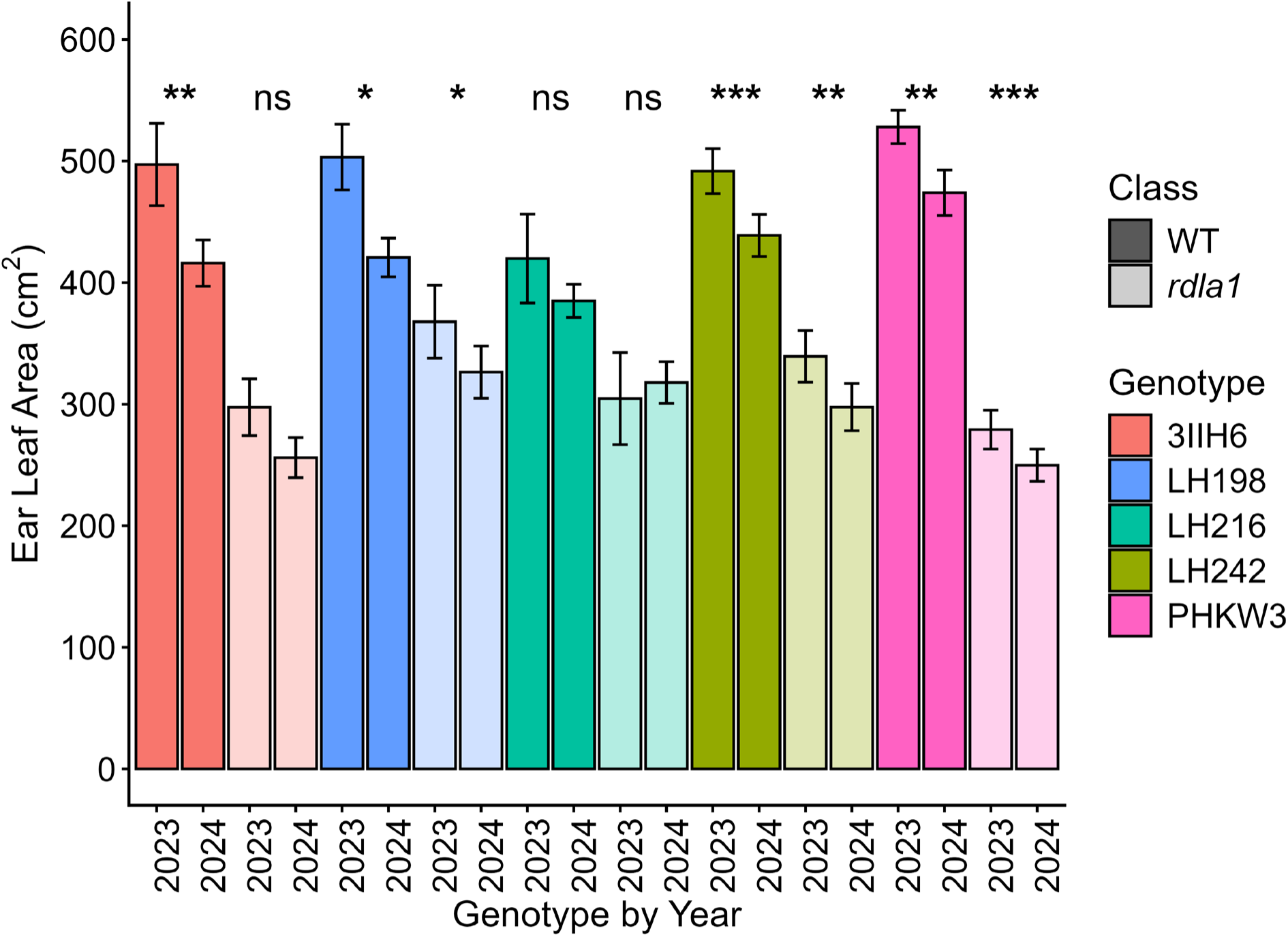
Absolute ear leaf area reduction ± standard error of *rdla1* introgression lines and wild-type genotypes that were replicated across both field trial seasons. Each bar represents the average of n = 4 plots (except 3IIH6 2023 is n = 2 plots). Bar colors represent the genotypic background evaluated. Bars with solid coloring are the wild-type version of each genotype, while bars with faded coloring are the *rdla1* version of each genotype. Significant differences calculated using a two-tailed Welch’s *t*-test are denoted as follows: *** = p < 0.001, ** = p < 0.01, * = p < 0.05, ns = nonsignificant.

**Supplementary Figure 7:**
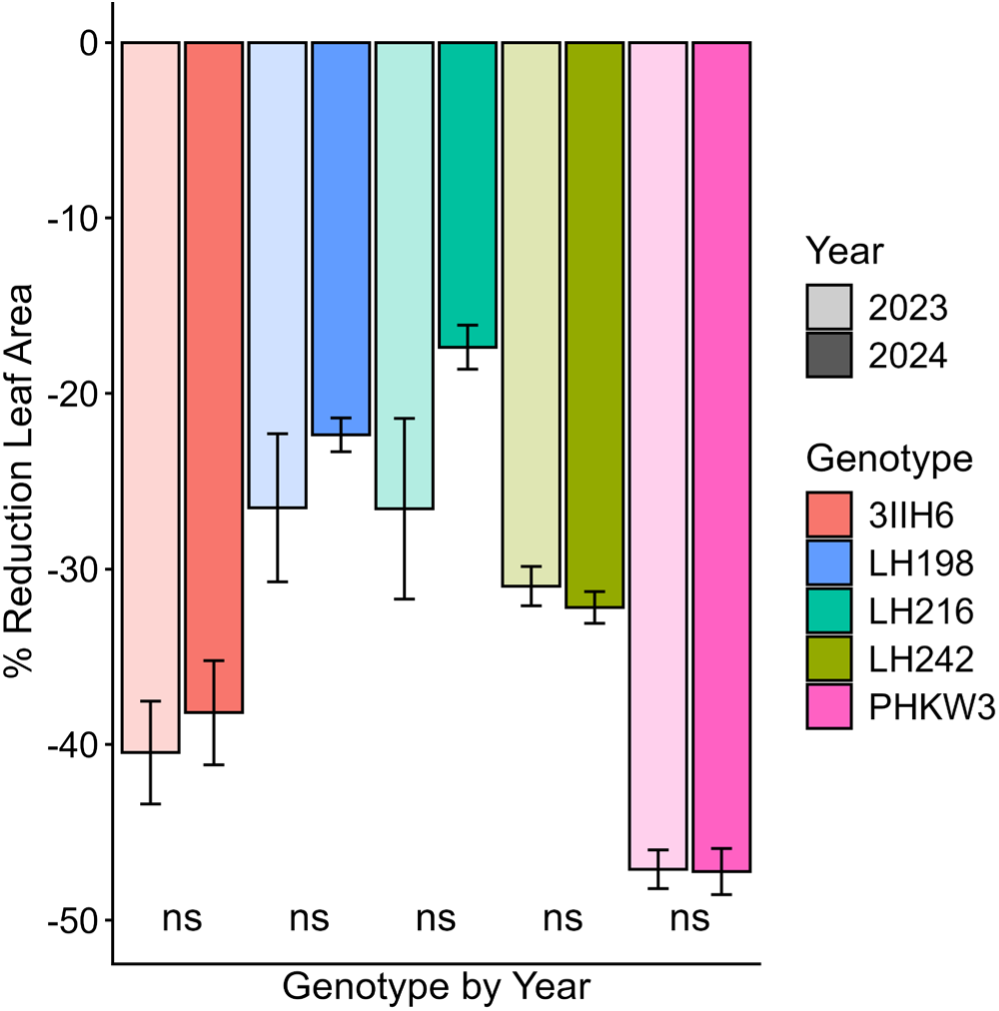
Relative ear leaf area reduction ± standard error of *rdla1* introgression lines relative to wild-type for genotypes that were replicated across both field trial seasons. Each bar represents the average of n = 4 plots (except 3IIH6 2023 is n = 2 plots). Bar colors represent the genotypic background evaluated. Bars with faded coloring are values for the 2023 field season, while bars solid coloring are values for the 2024 field season. Significant differences calculated using a two-tailed Welch’s *t*-test are denoted as follows: *** = p < 0.001, ** = p < 0.01, * = p < 0.05, ns = nonsignificant.

**Supplementary Figure 8:**
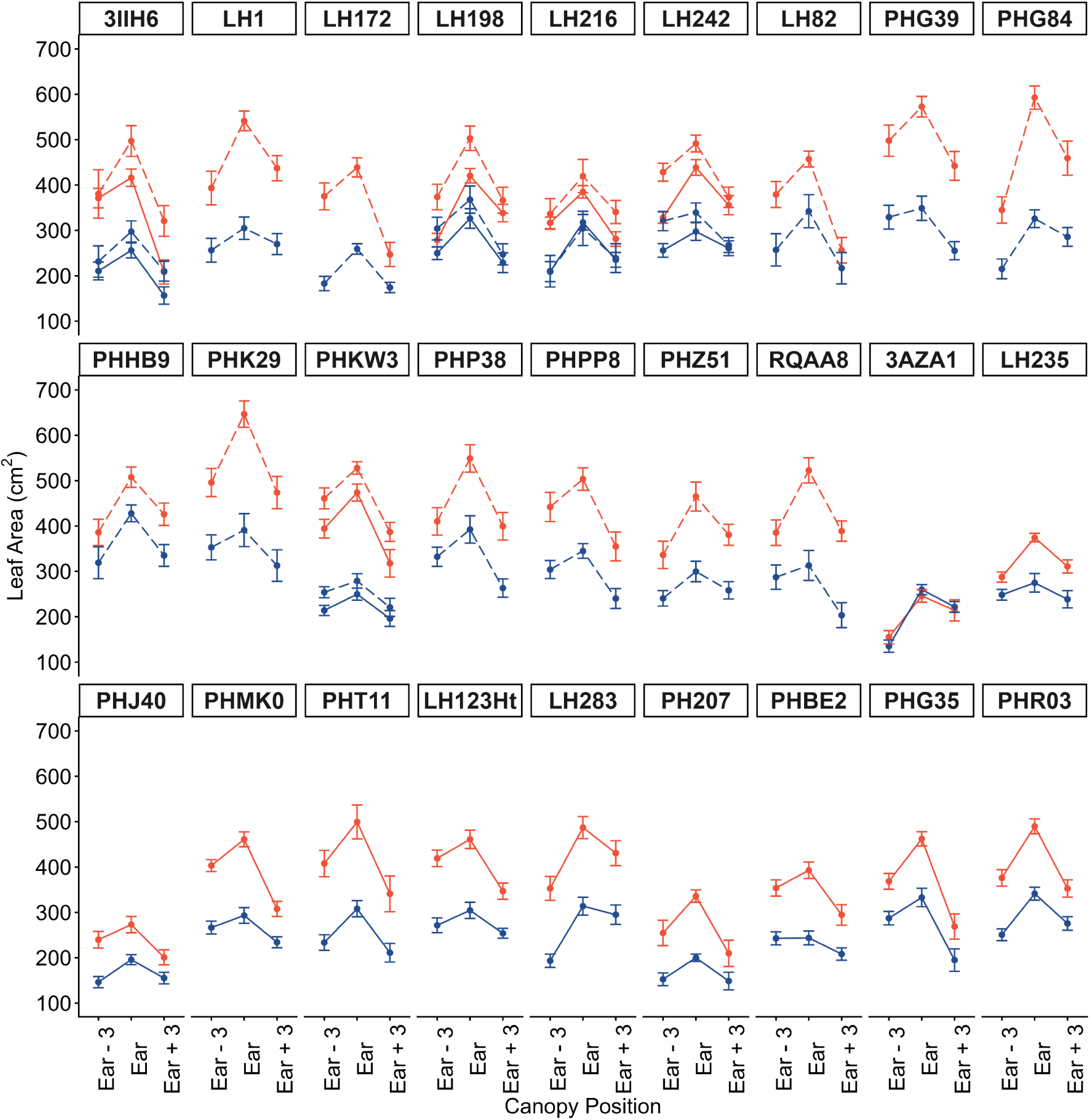
Average absolute leaf area reduction ± standard error of *rdla1* introgression lines and wild-type at the third leaf below the ear-bearing node, the leaf of the ear-bearing node, and the third leaf above the ear bearing node. Each bar represents the average of n = 4 plots (except 3IIH6 2023 is n = 2 plots). Orange points are wild-type values, and blue points are *rdla1* values. Genotypes with dashed lines were grown in the 2023 field trial, while genotypes with solid lines were grown during the 2024 field trial.

**Supplementary Figure 9:**
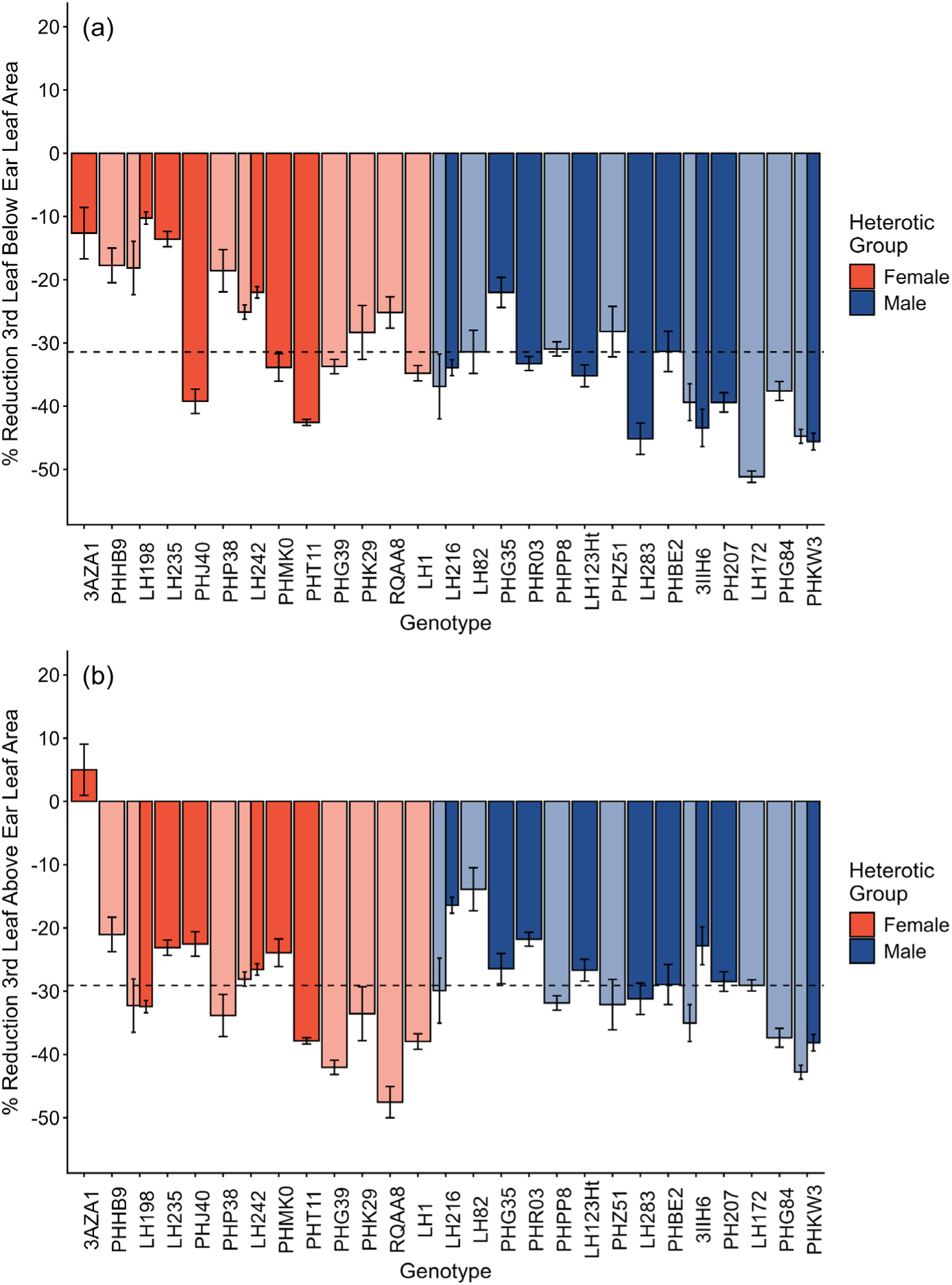
Average percentage leaf area reduction ± standard error of *rdla1* introgression lines relative to wild-type at the third leaf below the ear-bearing node (a) and the third leaf above the ear-bearing node (b). Each bar represents the average of n = 4 plots (except 3IIH6 2023 is n = 2 plots). Orange bars are inbreds classified as female, and blue bars are inbreds classified as male. Genotypes with faded coloring were grown in the 2023 field trial, while genotypes with solid coloring were grown during the 2024 field trial.

## References

1 Lee, E. A., & Tollenaar, M. (2007). Physiological basis of successful breeding strategies for maize grain yield. Crop Science, 47, S-202.

2 Monteith, J. L. (1972). Solar radiation and productivity in tropical ecosystems. Journal of Applied Ecology, 9(3), 747–766.

3 Long, S. P., Zhu, X. G., Naidu, S. L., & Ort, D. R. (2006). Can improvement in photosynthesis increase crop yields? Plant, Cell & Environment, 29(3), 315–330.

4 Kalogeropoulos, G., Elli, E. F., Trifunovic, S., & Archontoulis, S. V. (2024). Historical increases of maize leaf area index in the US Corn Belt due primarily to plant density increases. Field Crops Research, 318, 109615.

5 Perry, E. D., Hennessy, D. A., & Moschini, G. (2022). Uncertainty and learning in a technologically dynamic industry: Seed density in US maize. American Journal of Agricultural Economics, 104(4), 1388–1410.

6 Duvick, D. N. (2005). The contribution of breeding to yield advances in maize (Zea mays L.). Advances in Agronomy, 86, 83–145.

7 Gonzalez, V. H., Tollenaar, M., Bowman, A., Good, B., & Lee, E. A. (2018). Maize yield potential and density tolerance. Crop Science, 58(2), 472–485.

8 Assefa, Y., Carter, P., Hinds, M., Bhalla, G., Schon, R., Jeschke, M., … & Ciampitti, I. A. (2018). Analysis of long term study indicates both agronomic optimal plant density and increase maize yield per plant contributed to yield gain. Scientific Reports, 8(1), 4937.

9 Testa, G., Reyneri, A., & Blandino, M. (2016). Maize grain yield enhancement through high plant density cultivation with different inter-row and intra-row spacings. European Journal of Agronomy, 72, 28–37.

10 Maddonni, G. A., & Otegui, M. E. (2004). Intra-specific competition in maize: early establishment of hierarchies among plants affects final kernel set. Field Crops Research, 85(1), 1–13.

11 Haegele, J. W., Becker, R. J., Henninger, A. S., & Below, F. E. (2014). Row Arrangement, Phosphorus Fertility, and Hybrid Contributions to Managing Increased Plant Density of Maize. Agronomy Journal, 106, 1838–1846.

12 Maddonni, G. A., Otegui, M. E., & Cirilo, A. G. (2001). Plant population density, row spacing and hybrid effects on maize canopy architecture and light attenuation. Field Crops Research, 71(3), 183–193.

13 Duvick, D. N., & Cassman, K. G. (1999). Post–green revolution trends in yield potential of temperate maize in the North-Central United States. Crop Science, 39(6), 1622–1630.

14 Li, Y., Ma, X., Wang, T., Li, Y., Liu, C., Liu, Z., … & Smith, S. (2011). Increasing maize productivity in China by planting hybrids with germplasm that responds favorably to higher planting densities. Crop Science, 51(6), 2391–2400.

15 Mansfield, B. D., & Mumm, R. H. (2014). Survey of plant density tolerance in US maize germplasm. Crop Science, 54(1), 157–173.

16 Lambert, R. J., Mansfield, B. D., & Mumm, R. H. (2014). Effect of leaf area on maize productivity. Maydica, 59 (2014), 58–63.

17 Sangoi, L., Gracietti, M. A., Rampazzo, C., & Bianchetti, P. (2002). Response of Brazilian maize hybrids from different eras to changes in plant density. Field Crops Research, 79(1), 39–51.

18 Tian, F., Bradbury, P. J., Brown, P. J., Hung, H., Sun, Q., Flint-Garcia, S., … & Buckler, E. S. (2011). Genome-wide association study of leaf architecture in the maize nested association mapping population. Nature Genetics, 43(2), 159–162.

19 Ku, L., Ren, Z., Chen, X., Shi, Y., Qi, J., Su, H., … & Chen, Y. (2016). Genetic analysis of leaf morphology underlying the plant density response by QTL mapping in maize (Zea mays L.). Molecular Breeding, 36, 1–16.

20 Perez, R. P., Fournier, C., Cabrera-Bosquet, L., Artzet, S., Pradal, C., Brichet, N., … & Tardieu, F. (2019). Changes in the vertical distribution of leaf area enhanced light interception efficiency in maize over generations of selection. Plant, Cell & Environment, 42(7), 2105–2119.

21 Lambert, R. J., & Johnson, R. R. (1978). Leaf angle, tassel morphology, and the performance of maize hybrids 1. Crop Science, 18(3), 499–502.

22 Moon, J., Candela, H., & Hake, S. (2013). The Liguleless narrow mutation affects proximal-distal signaling and leaf growth. Development, 140(2), 405–412.

23 Buescher, E. M., Moon, J., Runkel, A., Hake, S., & Dilkes, B. P. (2014). Natural variation at sympathy for the ligule controls penetrance of the semidominant Liguleless narrow-R mutation in Zea mays. G3: Genes, Genomes, Genetics, 4(12), 2297–2306.

24 Scanlon, M. J., Schneeberger, R. G., & Freeling, M. (1996). The maize mutant narrow sheath fails to establish leaf margin identity in a meristematic domain. Development, 122(6), 1683–1691.

25 Scanlon, M. J. (2000). NARROW SHEATH1 functions from two meristematic foci during founder-cell recruitment in maize leaf development. Development, 127(21), 4573–4585.

26 Nardmann, J., Ji, J., Werr, W., & Scanlon, M. J. (2004). The maize duplicate genes narrow sheath1 and narrow sheath2 encode a conserved homeobox gene function in a lateral domain of shoot apical meristems.

27 Schneeberger, R., Tsiantis, M., Freeling, M., & Langdale, J. A. (1998). The rough sheath2 gene negatively regulates homeobox gene expression during maize leaf development. Development, 125(15), 2857–2865.

28 Timmermans, M. C., Hudson, A., Becraft, P. W., & Nelson, T. (1999). ROUGH SHEATH2: a Myb protein that represses knox homeobox genes in maize lateral organ primordia. Science, 151–153.

29 Tsiantis, M., Schneeberger, R., Golz, J. F., Freeling, M., & Langdale, J. A. (1999). The maize rough sheath2 gene and leaf development programs in monocot and dicot plants. Science, 284(5411), 154–156.

30 Lambert, R. J. (2010). Divergent selection for ear leaf area in maize. Maydica, 55(2), 155–161.

31 Jenks, M. A., Tuttle, H. A., Eigenbrode, S. D., & Feldmann, K. A. (1995). Leaf epicuticular waxes of the eceriferum mutants in Arabidopsis. Plant Physiology, 108(1), 369.

32 Rashotte, A. M., Jenks, M. A., & Feldmann, K. A. (2001). Cuticular waxes on eceriferum mutants of Arabidopsis thaliana. Phytochemistry, 57(1), 115–123.

33 Post-Beittenmiller, D. (1998). The cloned Eceriferum genes of Arabidopsis and the corresponding Glossy genes in maize. Plant Physiology and Biochemistry, 36(1-2), 157–166.

34 Zheng, J., He, C., Qin, Y., Lin, G., Park, W. D., Sun, M., … & Liu, S. (2019). Co-expression analysis aids in the identification of genes in the cuticular wax pathway in maize. The Plant Journal, 97(3), 530–542.

35 Jenks, M. A., Rashotte, A. M., Tuttle, H. A., & Feldmann, K. A. (1996). Mutants in Arabidopsis thaliana altered in epicuticular wax and leaf morphology. Plant Physiology, 110(2), 377–385.

36 Guo, T., Wang, D., Fang, J., Zhao, J., Yuan, S., Xiao, L., & Li, X. (2019). Mutations in the rice OsCHR4 gene, encoding a CHD3 family chromatin remodeler, induce narrow and rolled leaves with increased cuticular wax. International Journal of Molecular Sciences, 20(10), 2567.

37 Zhao, Y., Liu, Q., Wang, X., Zhang, W., Xu, W., Zhang, Y., & Liu, B. (2024). ZmCER1, a putative ECERIFERUM 1 protein in maize, functions in cuticular wax biosynthesis and bulliform cell development. The Crop Journal, 12(3), 743–752.

38 Bourdenx, B., Bernard, A., Domergue, F., Pascal, S., Léger, A., Roby, D., … & Joubès, J. (2011). Overexpression of Arabidopsis ECERIFERUM1 promotes wax very-long-chain alkane biosynthesis and influences plant response to biotic and abiotic stresses. Plant physiology, 156(1), 29–45.

39 Grünhofer, P., Herzig, L., Zhang, Q., Vitt, S., Stöcker, T., Malkowsky, Y., … & Schreiber, L. (2024). Changes in wax composition but not amount enhance cuticular transpiration. Plant, Cell & Environment, 47(1), 91–105.

40 Shellakkutti, N., Thangamani, P. D., Suresh, K., Baales, J., Zeisler-Diehl, V., Klaus, A., … & Kreszies, T. (2022). Cuticular transpiration is not affected by enhanced wax and cutin amounts in response to osmotic stress in barley. Physiologia Plantarum, 174(4), e13735.

41 Ku, L. X., Zhang, J., Guo, S. L., Liu, H. Y., Zhao, R. F., & Chen, Y. H. (2012). Integrated multiple population analysis of leaf architecture traits in maize (Zea mays L.). Journal of Experimental Botany, 63(1), 261–274.

42 Wang, H., Liang, Q., Li, K., Hu, X., Wu, Y., Wang, H., … & Huang, C. (2017). QTL analysis of ear leaf traits in maize (Zea mays L.) under different planting densities. The Crop Journal, 5(5), 387–395.

43 Cui, T., He, K., Chang, L., Zhang, X., Xue, J., Liu, J. (2017). QTL mapping for leaf area in maize (Zea mays L.) under multi-environments. Journal of Integrative Agriculture, 16(4), 800–808.

44 Khaipho-Burch, M., Cooper, M., Crossa, J., de Leon, N., Holland, J., Lewis, R., … & Buckler, E. S. (2023). Genetic modification can improve crop yields—but stop overselling it. Nature, 621(7979), 470–473.

45 Fulton, T. M., Chunwongse, J., & Tanksley, S. D. (1995). Microprep protocol for extraction of DNA from tomato and other herbaceous plants. Plant Molectular Biology Reporter, 13(3), 207–209.

46 Clark, L. V., Brummer, J. E., Głowacka, K., Hall, M. C., Heo, K., Peng, J., … & Sacks, E. J. (2014). A footprint of past climate change on the diversity and population structure of Miscanthus sinensis. Annals of Botany, 114(1), 97–107.

47 Poland, J. A., & Rife, T. W. (2012). Genotyping-by-sequencing for plant breeding and genetics. The Plant Genome, 5(3).

48 Catchen, J., Hohenlohe, P. A., Bassham, S., Amores, A., & Cresko, W. A. (2013). Stacks: an analysis tool set for population genomics. Molecular Ecology, 22(11), 3124–3140.

49 Langmead, B., & Salzberg, S. L. (2012). Fast gapped-read alignment with Bowtie 2. Nature Methods, 9(4), 357–359.

50 Danecek, P., Schiffels, S., & Durbin, R. (2014). Multiallelic calling model in bcftools (-m). https://samtools.github.io/bcftools/call-m.pdf.

51 Danecek, P., Auton, A., Abecasis, G., Albers, C. A., Banks, E., DePristo, M. A., … & 1000 Genomes Project Analysis Group. (2011). The variant call format and VCFtools. Bioinformatics, 27(15), 2156–2158.

52 Afgan, E., Baker, D., Batut, B., Van Den Beek, M., Bouvier, D., Čech, M., … & Blankenberg, D. (2018). The Galaxy platform for accessible, reproducible and collaborative biomedical analyses: 2018 update. Nucleic Acids Research, 46(W1), W537–W544.

53 Cingolani, P., Platts, A., Wang, L. L., Coon, M., Nguyen, T., Wang, L., … & Ruden, D. M. (2012). A program for annotating and predicting the effects of single nucleotide polymorphisms, SnpEff: SNPs in the genome of Drosophila melanogaster strain w1118; iso-2; iso-3. Fly, 6(2), 80–92.

54 Cheng, H., Concepcion, G. T., Feng, X., Zhang, H., & Li, H. (2021). Haplotype-resolved de novo assembly using phased assembly graphs with hifiasm. Nature Methods, 18(2), 170–175.

55 Ye, J., Coulouris, G., Zaretskaya, I., Cutcutache, I., Rozen, S., & Madden, T. L. (2012). Primer-BLAST: a tool to design target-specific primers for polymerase chain reaction. BMC Bioinformatics, 13(1), 134.

56 Ruzin, S. E. (1999). Plant microtechnique and microscopy. New York: Oxford University Press.

57 Johansen, D. A. (1940). Plant Microtechnique, 1st Edition. McGraw-Hill Book Co. Ltd. New York.

58 Sylvester, A. W., & Ruzin, S. E. (1994). Light microscopy I: dissection and microtechnique. In The Maize Handbook (pp. 83–95). New York, NY: Springer New York.

59 Zhang, J. Y., Broeckling, C. D., Sumner, L. W., & Wang, Z. Y. (2007). Heterologous expression of two Medicago truncatula putative ERF transcription factor genes, WXP1 and WXP2, in Arabidopsis led to increased leaf wax accumulation and improved drought tolerance, but differential response in freezing tolerance. Plant Molecular Biology, 64(3), 265–278.

60 R Core Team (2025). R: A Language and Environment for Statistical Computing. R Foundation for Statistical Computing, Vienna, Austria. URL https://www.R-project.org/.

61 Posit Team (2025). RStudio: Integrated Development Environment for R. Posit Software, PBC, Boston, MA. URL http://www.posit.co/.

